# Connexin43 in mesenchymal lineage cells regulates body adiposity and energy metabolism in mice

**DOI:** 10.1101/2024.01.05.574415

**Authors:** Seung-Yon Lee, Francesca Fontana, Toshifumi Sugatani, Ignacio Portales Castillo, Giulia Leanza, Ariella Coler-Reilly, Roberto Civitelli

## Abstract

Connexin43 (Cx43) is the most abundant gap junction protein present in the mesenchymal lineage. In mature adipocytes, Cx43 mediates white adipose tissue (WAT) “beiging” in response to cold exposure and maintains the mitochondrial integrity of brown adipose tissue (BAT). We found that genetic deletion of *Gja1* (Cx43 gene) in cells that give rise to chondro-osteogenic and adipogenic precursors driven by the *Dermo1/Twist2* promoter leads to lower body adiposity and partial protection against the weight gain and metabolic syndrome induced by a high fat diet (HFD) in both sexes. These protective effects from obesogenic diet are related to increased locomotion, fuel utilization, energy expenditure, non-shivering thermogenesis, and better glucose tolerance in conditionally *Gja1* ablated mice. Accordingly, *Gja1* mutant mice exhibit reduced adipocyte hypertrophy, partially preserved insulin sensitivity, increased BAT lipolysis and decreased whitening under HFD. This metabolic phenotype is not reproduced with more restricted *Gja1* ablation in differentiated adipocytes, suggesting that Cx43 has a hitherto unknown function in adipocyte progenitors or other targeted cells, resulting in restrained energy expenditures and fat accumulation. These results disclose an hitherto unknown action of Cx43 in adiposity, and offer a promising new pharmacologic target for improving metabolic balance in diabetes and obesity.

**Graphical Abstract:** 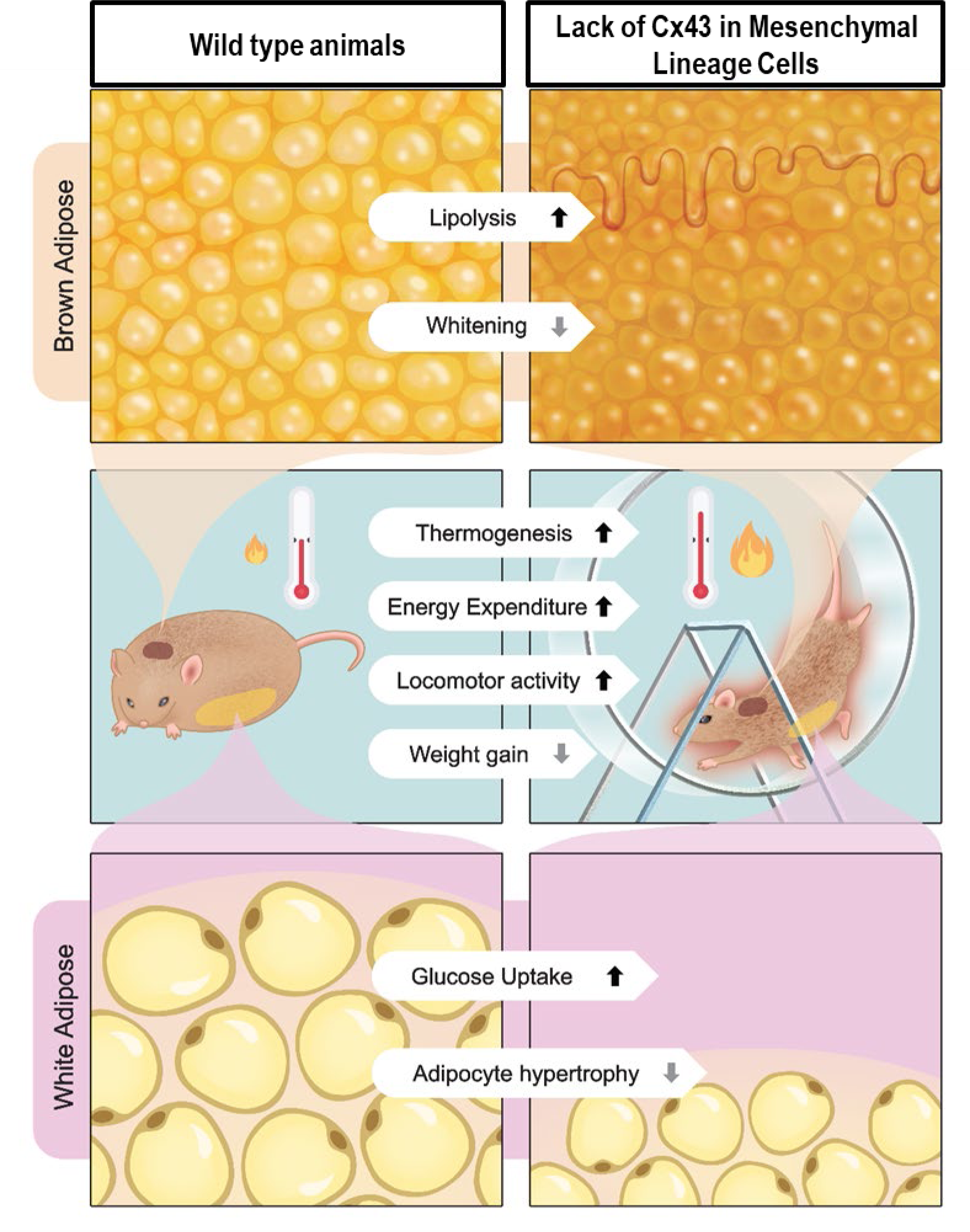

## Introduction

The adipose tissue is a metabolically active organ involved in many aspects of normal physiology, including tissue nutrient homeostasis and energy metabolism (1). The size of an individual’s adipose tissues depends on developmental and environmental factors that affect the number and size of adipocytes. Indeed, body adiposity can increase by either volumetric expansion of existing adipocytes (hypertrophy), or increased number of adipocytes (hyperplasia). The balance between hypertrophic expansion of existing adipocyte and adipogenic differentiation is an integral component of metabolism and affects an individual’s health (2).

White adipocytes make up the bulk of fatty tissue in most animals, their main role being long-term energy storage in fat depots, which are referred to as white adipose tissue (WAT). In conditions of increased energy needs, white adipocytes can activate lipolysis and release free fatty acids from triglycerides. The WAT also has an endocrine function, producing leptin, adiponectin and other adipokines that regulate fuel utilization in peripheral tissues and lipid metabolism (3). Brown and beige adipocytes are instead primarily involved in energy consumption and heat dissipation. Brown adipocytes are found in a specific depot, the brown adipose tissue (BAT), while beige adipocytes are present in WAT under conditions of increased energy demands. The BAT contributes to lowering circulating lipids by favoring triglyceride uptake via lipoprotein lipase (4), fatty acid uptake from triglyceride-rich lipoproteins, and enhanced hepatic clearance of cholesterol-enriched molecules (5). Thus, the adipose tissue functions to balance an individual’s needs for energy storage, utilization, and dissipation as heat.

Our work explores the role of a gap junction protein in adipose tissue function. Gap junctions are intercellular channels that allow aqueous continuity between connected cells. They are formed by juxtaposition of two transmembrane “hemichannels” in apposing cells, with each hemichannel composed of a homo- or hetero-hexamer of connexin proteins. Gap junctions have been observed in adipose tissue during development (6, 7) and BAT adipocytes are more functionally connected than white adipocytes in adult mice (8). However, the specific function of gap junctions in the different adipose tissues remains to be fully elucidated. Among the 21 connexins present in humans (9), connexin43 (Cx43) is the main gap junction protein expressed in mesenchymal lineage cells (9, 10). Work using adipocyte-targeted *Gja1* (Cx43 gene) ablation in mice shows that Cx43 mediates adipocyte beiging in response to sympathetic neuronal signals, and that overexpression of Cx43 promotes WAT beiging even with mild cold stimuli (11). Moreover, *Gja1* deletion in mature adipocytes disrupts mitochondrial integrity via autophagy resulting in metabolic dysfunction in mice kept on a high-fat diet (12). These results suggest that Cx43 in mature adipocytes has beneficial roles in controlling energy metabolism and thermogenesis.

In our studies on Cx43 in the skeleton, we had observed that mice with ablation of *Gja1* broadly in mesenchymal lineage cells, using the *Dermo-1/Twist-2* promoter, acquire less body weight with age, in addition to having undermineralized, thinner long bones relative to control mice (13). Using these genetically engineered mice, here we show that, contrary to expectations, lack of Cx43 in mesenchymal lineage cells is protective against the metabolic stress produced by a high calorie diet, whereas more restricted *Gja1* ablation in mature adipocytes is not. Specifically, we demonstrate that lack of Cx43 reduces white adipocyte hypertrophy, increases glucose uptake and insulin sensitivity, prevents BAT whitening, and increases locomotor activity, energy expenditure, non-shivering thermogenesis, and BAT lipolysis. Thus, Cx43 has complex and cell-specific functions in adipose tissue biology and energy metabolism.

## Results

### *Gja1* ablation in mesenchymal lineage cells results in metabolic benefits in mice on standard diet

To study the role of Cx43 in mesenchymal progenitors, we used *Dermo1/Twist2-Cre:Gja1^flox/flox^*(cKO^Tw2^) mice, which we have previously described (13, 14). *Dermo1/Twist2-Cre* (Tw2-Cre) targets the entire skeleton, muscle, and epidermis at birth (Fig. S1A). In adult *Dermo1/Twist2-Cre;Ai9* reporter mice, TdT signal was detected diffusely in BAT and WAT adipocytes, including cells in the stromal/perivascular areas, although vessels were not targeted (Fig. S1B, C). A small proportion of hepatocytes were also targeted by *Dermo1/Twist2-Cre* (Fig. S1D), and in the spleen, TdT fluorescence was abundant in the stromal component while the nodular areas were negative (Fig. S1E). Abundant X-gal staining of fat depots and in histologic sections of BAT and WAT confirmed effective Tw2-Cre-driven *Gja1* gene recombination in both fat tissues (Fig. S1 F-I). The more intense stain in BAT (Fig. S1F’, G’) most likely reflects higher cell density than in WAT (Fig. S1H’, I’).

At age 7 months, body weight was marginally lower in cKO^Tw2^ than in *Gja1^flox/flox^* (wild type equivalent: WT) male mice (Fig. 1A), and the difference was more pronounced among female mice (Fig. S2A). Neither fat mass nor lean mass were altered in mutant mice when assessed by dual-energy x-ray absorptiometry (Fig. 1B, C; S2B, C). At necropsy, gonadal fat depots (Fig. 1D; S2D) and BAT mass were similar in control and mutant mice (Fig. 1E; S2E). However, the weight of inguinal fat depots from 7-months-old cKO^Tw2^ mice was lower than those excised from WT mice of both sexes (Fig. 1D; S2F). Histologically, we observed no abnormalities in morphology or size of adipocytes in inguinal WAT, or in BAT in mutant male mice (Fig. 1F). Following an intraperitoneal glucose tolerance test (iGTT), male cKO^Tw2^ mice showed faster return to baseline of blood glucose compared to WT mice (Fig. 1G). Such differences were not seen in female cKO^Tw2^ mice, which instead were hyperglycemic after glucose load (Fig. S2G, H). Indirect calorimetry showed no significant differences between genotypes in food consumption (Fig. S3A, B, Table S1), energy expenditure (Fig. S3C, D, Table S1), and RER (Fig. S3E, F, Table S1). However, cKO^Tw2^ mice were more active than WT mice during dark hours (Fig. S3G, H, Table S1).

**Figure 1.**
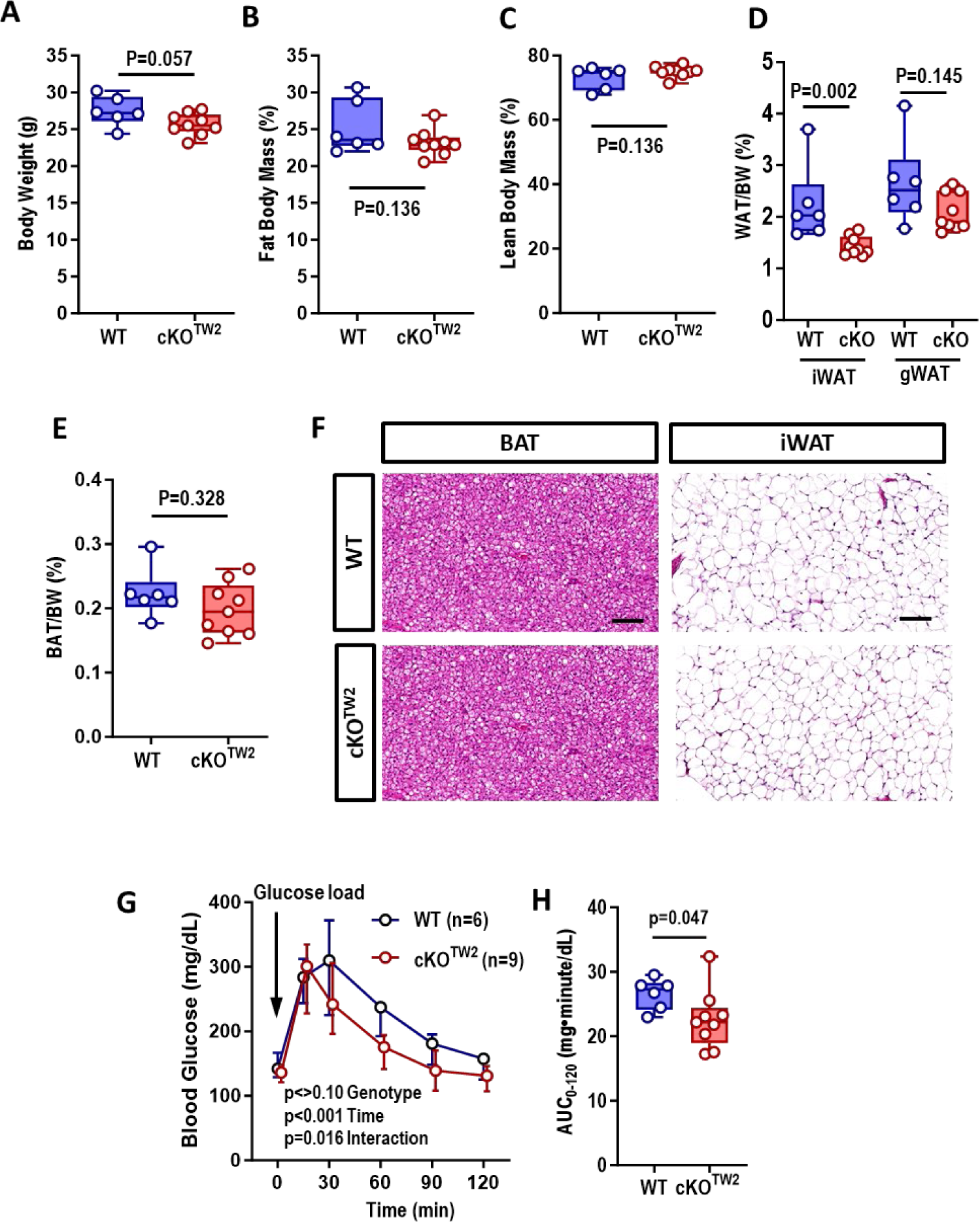
*Gja1* ablation in mesenchymal lineage cells leads to metabolic benefits in regular chow died (RCD) fed male mice on standard diet. (A) Body weight, and (B) DXA-determined percent body fat and (C) lean mass in 7-month-old wild type (WT, blue) and cKO^TW2^ mice (red). (D) White adipose tissue (WAT) weight normalized to body weight (BW) in inguinal (iWAT) and gonadal (gWAT) depots. (E) Weight of suprascapular brown adipose tissue (BAT) depots normalized to BW. (F) Hematoxylin-eosin-stained histologic sections of BAT and iWAT from wild type and cKO^TW2^ mice (bar=100 µm). Each image is representative of 6 wild type and 9 cKO^TW2^ mice. (G) Intraperitoneal glucose tolerance test: blood glucose before and after an intraperitoneal load of 1.5g/kg D-glucose. (H) Areas under the curve (AUC) calculated between 0 and 120 minutes for animals included in panel G. Data are presented as boxplots representing the interquartile range with median (inside bar). (A-E, H) Groups were compared using two-tailed Mann-Whitney U-test. (G) P-values represent the effect of genotype, time and their interaction by two-way ANOVA (genotype, F=3.082, p=0.103; time: F=97.14, p<0.001; time x genotype: F=3.032, p=0.016).

### Cx43 is up-regulated by high fat diet in BAT, while it is down-regulated during white adipocyte differentiation

Cx43 is the most abundant connexin in the adipose tissue (8); it is up-regulated during WAT beiging in mice (11), and during adipogenesis in human induced pluripotent stem cells (15). Using RT-qPCR, we confirmed expression of *Gja1* in BAT, and up-regulation in mice fed a high fat diet (HFD), with fat providing 60% of calorie intake (Fig. 2A). Of the other two connexins present in osteo-chondroprogenitor cells, *Gjc1* was also expressed in BAT and upregulated by HFD, even though it was about 10-fold less abundant than *Gja1* mRNA; whereas *Gja4* was not detected (Fig. 2A). Although mRNA for *Gja1* and *Gjc1* was barely detectable in WAT extracts regardless of the diet (Fig. 2A), Cx43 protein was abundantly expressed in pre-adipocytes obtained from the stromal-vascular fraction of WAT and in the IngWAT pre-adipocyte cell line (16) (Fig. 2B). Consistent with the mRNA results, Cx45 was also present in pre-adipocytes while Cx40 was undetectable; notably, both Cx43 and Cx45 protein and mRNA were rapidly downregulated upon in vitro adipocyte differentiation (Fig. 2C, D). Thus, low expression of Cx43 in differentiated adipocytes may explain the barely detectable *Gja1* mRNA in WAT extracts. Immunohistochemistry confirmed more intense Cx43-specific signal in BAT of HFD mice relative to mice kept on regular diet, and only background signal in BAT of cKO^Tw2^ mice (Fig. 2E, upper panels). Of note, Cx43-specific stain was absent in blood vessels, while it was visible in perivascular cells. Similarly, Cx43-specific stain was detected in WAT adipocytes kept on RCD, without notable increase in mice on HFD; whereas no Cx43-specific staining was detected in WAT of cKO^Tw2^ mice (Fig. 2E, lower panels).

**Figure 2.**
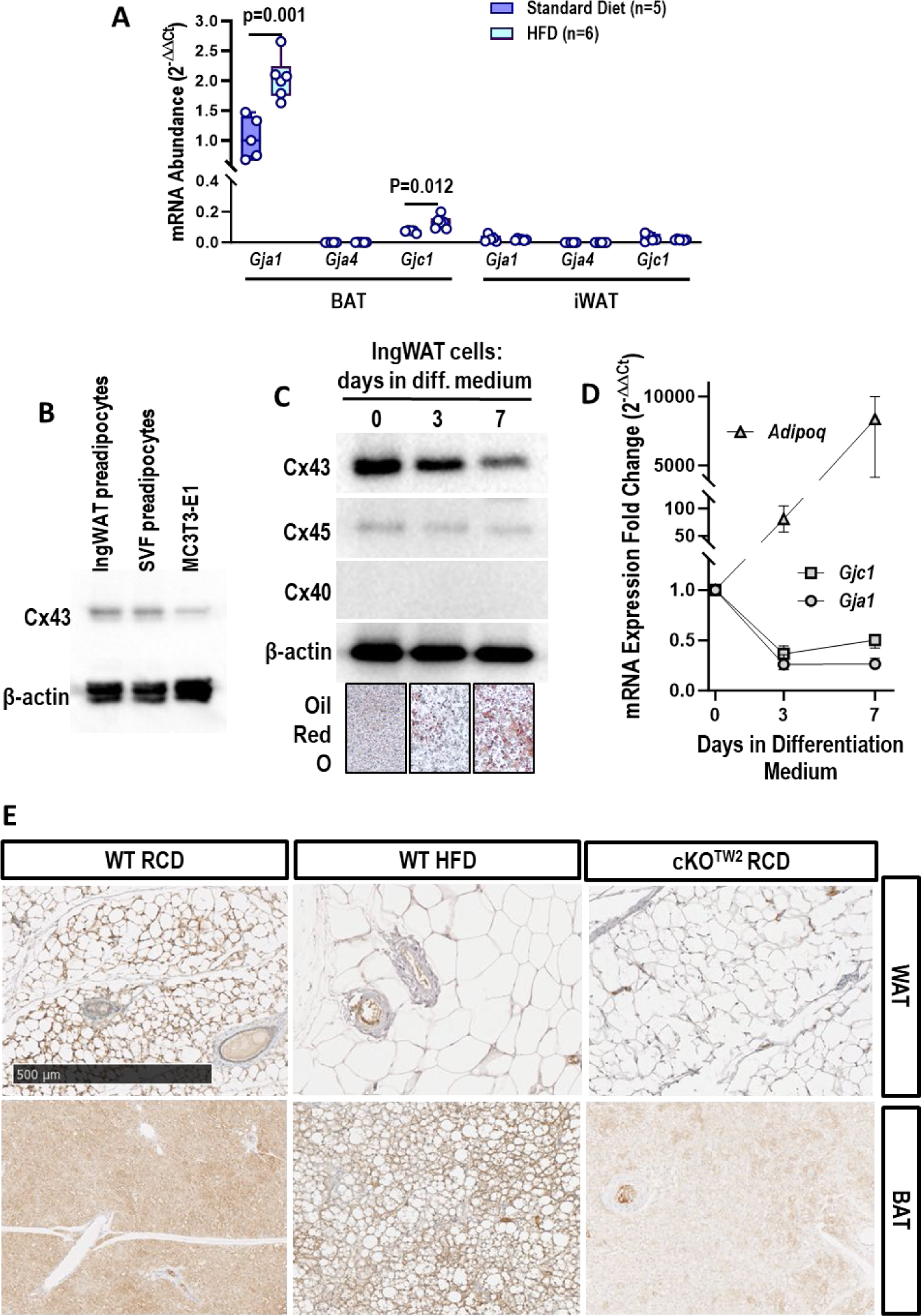
Cx43 is up-regulated by high fat diet (HFD) in brown adipose tissue (BAT), and down-regulated during adipogenic differentiation. (A) Expression of connexin mRNA by qPCR in inguinal white adipose tissue (iWAT and BAT) in 5-month-old male mice fed either a regular chow diet (RCD) or HFD for 12 weeks. Data are presented as boxplots representing the interquartile range with median (inside bar). Groups were compared using two-tailed Mann-Whitney U-test. (B) Western blot of whole cell lysates of confluent, undifferentiated cultures of IngWAT pre-adipocytic cells, cells isolated from the stromal-vascular fraction (SVF) of WAT, and the osteogenic cell line, MC3T3-E1, as control. (C) Western blot of whole cell lysates and (D) qRT-PCR analysis of mRNA of IngWAT cells before and during adipogenic differentiation (median and range, n=6; p<0.001 vs. time 0 at all time points). (E) Immunohistochemisty stain (brown color) for Cx43 in WAT and BAT isolated from WT mice kept of either RCD or HFD and in cKO^TW2^ mice. Scale bar: 500 µm. Each image is representative of three mice per condition.

### *Gja1* ablation in mesenchymal lineage cells is partially protective against high fat diet-induced metabolic syndrome and fat accumulation

We then investigated the consequences of Cx43 deletion on the metabolic response to obesogenic diets. Male mice fed a HFD for 12 weeks almost doubled their body weight, while cKO^Tw2^ mice experienced a far lesser weight gain on the same HFD (p<0.001 for effect of diet by 2-way ANOVA; Fig. 3A). Moreover, percent body fat was significantly lower, and percent lean weight was significantly higher in cKO^Tw2^ compared with WT male mice after HFD (Fig. 3B, C). Both WT and cKO^Tw2^ mice were hyperglycemic after 12 weeks on HFD. However, iGTT revealed a more efficient response in cKO^Tw2^ relative to WT mice, which remained severely hyperglycemic even 120 minutes after the glucose load (Fig. 3D, E). When HFD fed mice were injected insulin i.p. (ITT), blood glucose was significantly lower at all time-points in cKO^Tw2^ mice compared with WT (Fig. 3F, G). Similarly, female cKO^Tw2^ mutants gained far less weight on HFD (Fig. S4A) and accumulated significantly less body fat (Fig. S4B), resulting in smaller peripheral fat depots (Fig. S4C, D). Glucose tolerance test showed prolonged hyperglycemia in WT females fed HFD, while cKO^Tw2^ had significantly lower blood glucose levels (Fig. S4E, F), thus reversing the trend seen in female mice kept on chow diet (see above); however, both mutant and WT females responded equally to an insulin bolus (Fig. S4G, H). These results are also consistent with previous evidence that female mice are partially protected from the metabolic effects of HFD relative to male mice (17).

**Figure 3.**
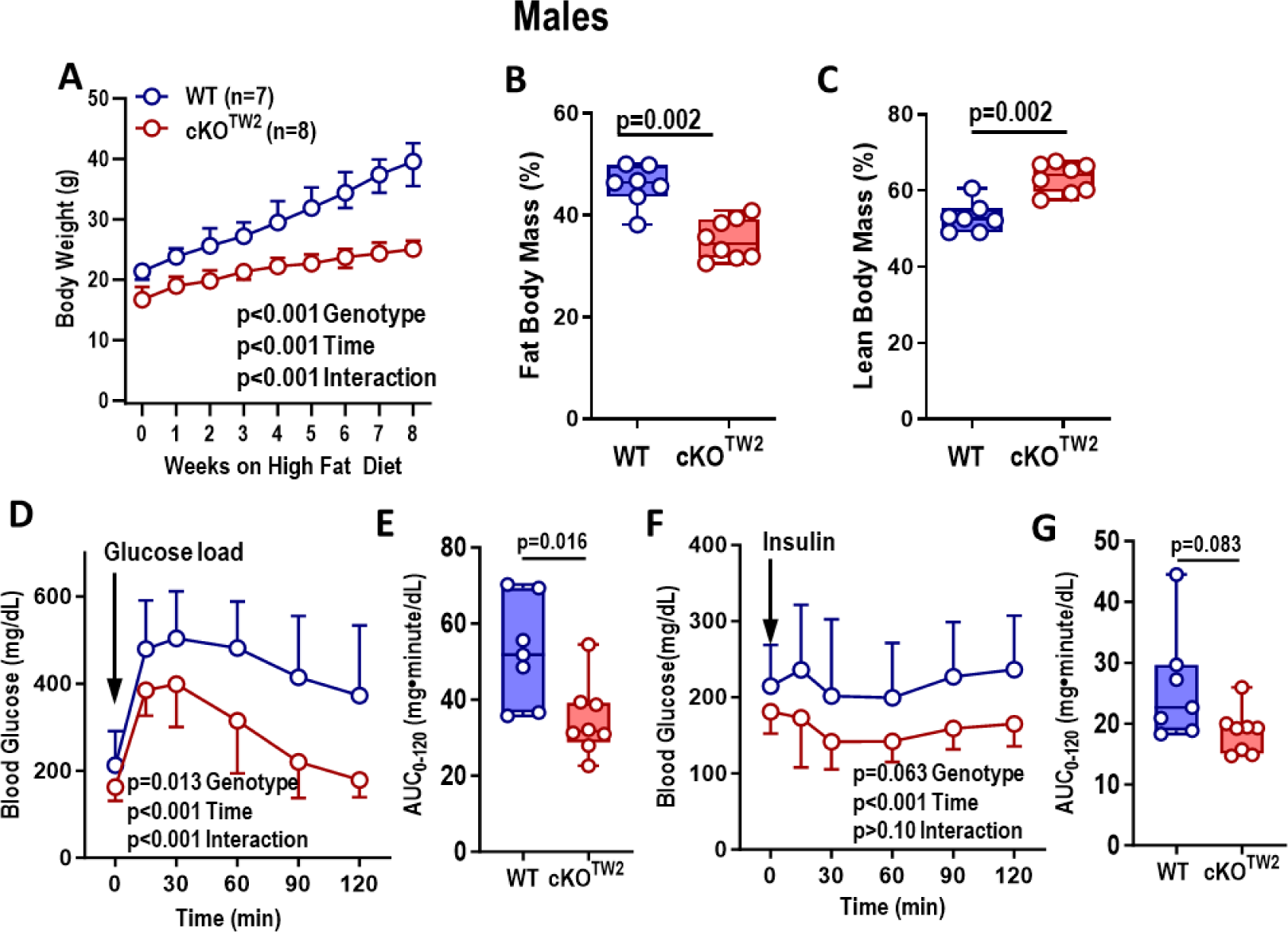
*Gja1* ablation in mesenchymal lineage cells is partially protective of high fat diet-induced obesity, hyperglycemia, and reduced glucose tolerance in male mice. (A) Body weight of 2-month-old wild type (WT, blue) and cKO^TW2^ (red) male mice during 8 weeks on HFD feeding. Data are shown as median and interquartile range; P-values represent the effect of genotype, time and their interaction by repeated measures two-way ANOVA (genotype, F=53.53, p<0.001; time: F=288.3, p<0.001; time x genotype: F=42.08, p<0.001). (B) Percent body fat and (C) lean mass by DXA after HFD in the two genotype groups. (D) Intraperitoneal glucose tolerance test: blood glucose before and after an intraperitoneal load of 1.5g/kg D-glucose (mean ± 95% CI; two-way ANOVA: genotype, F=8.202, p=0.013; time: F=70.87, p<0.001; time x genotype: F=5.972, p<0.001). (E) Areas under the curve (AUC) calculated between 0 and 120 minutes for animals included in panel D. (F) Intraperitoneal insulin tolerance test: blood glucose before and after an intraperitoneal injection of 0.75U/kg insulin (mean ± 95% CI; two-way ANOVA: genotype, F=4.131, p=0.063; time: F=6.634,p<0.001 ; time x genotype: F=1.347,p>0.10 ). (G) Areas under the curve (AUC) calculated between 0 and 120 minutes for animals included in panel F. Group data are in boxplots representing the interquartile range with median (inside bar), and P values were determined by two-sided Mann-Whitney U-test.

Given the more severe metabolic phenotype, the following studies were performed only in male mice. Consistent with the reduced obesity, WAT mass was macroscopically smaller in cKO^Tw2^ relative to WT male mice after 12 weeks on HFD (Fig. 4A). Expressed as percent body weight, both inguinal and gonadal WAT depots were comparatively smaller in cKO^Tw2^ than in WT mice (Fig. 4B). Importantly, H&E staining of iWAT sections revealed smaller sized adipocytes in cKO^Tw2^ mice, compared to larger cells in WT iWAT (Fig. 4C). As histologic sections of liver showed mild steatosis in WT mice but no fat accumulation in cKO^Tw2^ mutants following HFD (Fig. 4D), we can exclude a shift in fat accumulation into other organs in Cx43-deficient mice, as it occurs in lipodystrophy. Finally, serum insulin, C-peptide, Igf-1, and glucagon were all significantly lower in cKO^Tw2^ than in WT male mice (Fig. 4E-H), as were serum triglycerides, cholesterol, and free fatty acids (Fig. 4I-K). Overall, these results suggest that absence of Cx43 in the mesenchymal lineage is partially protective of the obesogenic effect and related metabolic abnormalities of high caloric dietary intake, a conclusion opposite to what has been reported when *Gja1* was ablated specifically in mature adipocytes (12).

**Figure 4.**
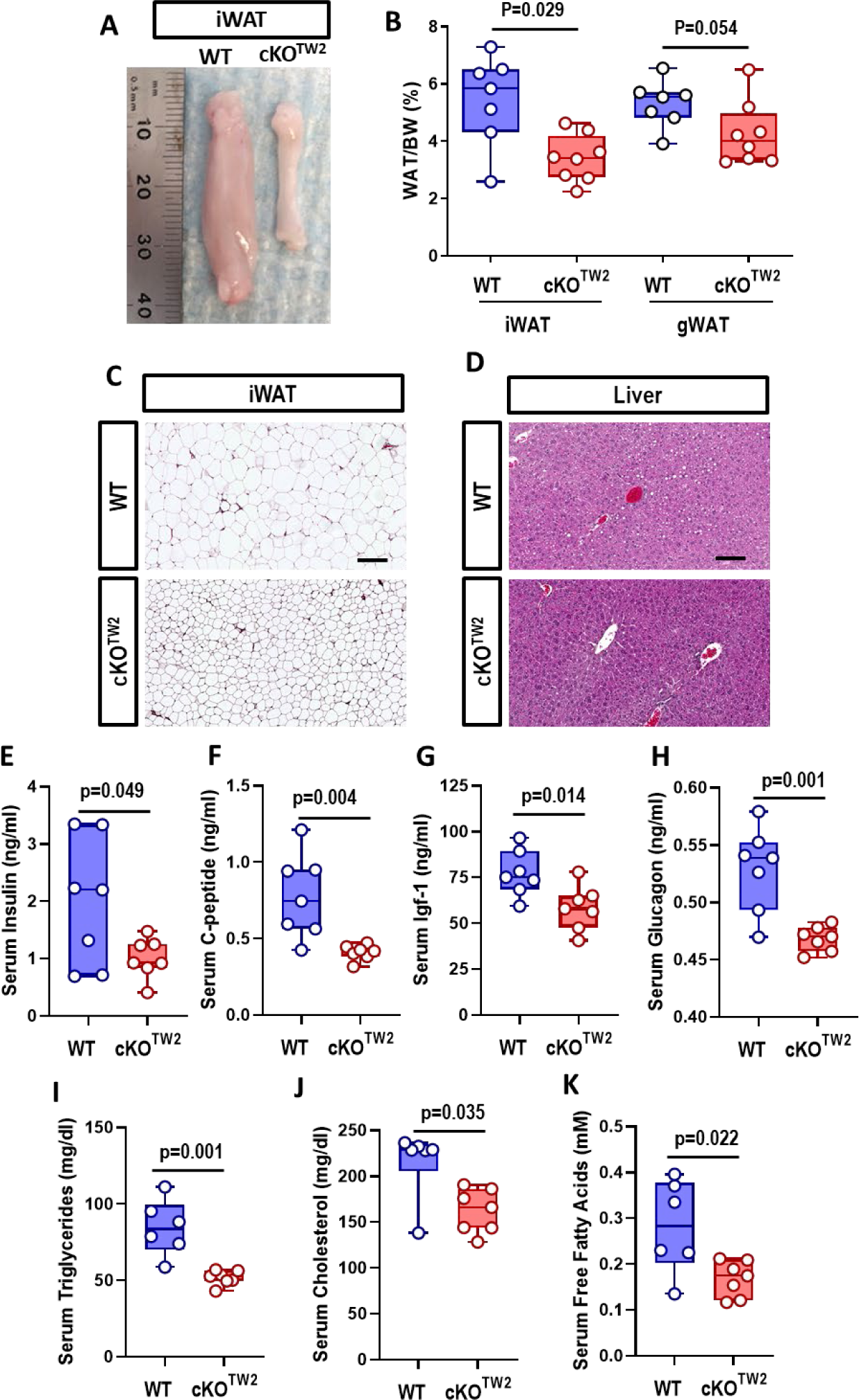
*Gja1* ablation in mesenchymal lineage cells protects high fat diet-induced expansion of fat depots, adipocyte hypertrophy, hyperinsulinemia, and hyperlipidemia in male mice. (A) Representative morphology of inguinal white adipose tissue (iWAT) in 2-month-old WT and cKO^TW2^ male mice after 12 weeks on a HFD. (B) Percent iWAT and gonadal WAT (gWAT) relative to body weight in the two genotypes after HFD (boxplots represent interquartile range with median); P values were determined by two-sided Mann-Whitney U-test. (C) Representative H&E-stained histological sections of iWAT and (D) liver of WT (n=7) and cKO^TW2^ (n=8) mice after 12 weeks on HFD. Scale bar: 50 µm. Serum levels of (E) insulin, (F) C-peptide, (G) Igf-1, (H) glucagon, (I) triglyceride, (J) cholesterol, and (K) fatty acid measured in WT and cKO^TW2^ mice after 12 weeks on HFD. Group data are in boxplots representing the interquartile range with median (inside bar), and P values were determined by two-sided Mann-Whitney U-test.

### *Gja1* ablation in mesenchymal lineage cells favors adipogenesis and glucose uptake in WAT of mice fed a high fat diet

To study the mechanisms of the partial resistance of Cx43-deficient mice to the obesogenic HFD, we examined the expression of genes associated with adipogenesis, glucose uptake, and lipolysis. After HFD, *Cebpα*, *Pparγ*, and *Adipoq*, were higher in subcutaneous fat (iWAT) obtained from cKO^Tw2^ relative to control mice, whereas *Lep* was lower (Fig. 5A). Expression of the two main glucose transporters, *Glut1* and *Glut4*, was also increased in iWAT of cKO^Tw2^ relative to WT mice (Fig. 5B). Basal glucose uptake, measured as fluorescence intensity in cells after incubation with 2-NBD-Glucose, was higher in adipocytes derived from cKO^Tw2^ relative to WT mice. Such difference was significant after 60 min incubation, and at 30 min in the presence of insulin (Fig. 5C). While insulin significantly stimulated glucose uptake in WT adipocytes at 60 min, it had no effect in cKO^Tw2^ adipocytes, in which basal glucose uptake at 60 min was similar to glucose uptake by WT adipocytes in the presence of insulin (Fig. 5C). These results suggest that basal glucose transport may be maximal in cKO^Tw2^ white adipocytes after high fat feeding. There were no differences in expression of lipolysis-related genes, lipoprotein lipase (*Lpl*) and adipose triglyceride lipase (*Pnpla2*) in the iWAT of mutant and WT mice (Fig. 5D). Likewise, glycerol release from iWAT cells, an index of lipolysis, was not different between genotypes either in basal conditions, or under β-adrenergic stimulus by isoproterenol (Fig. 5E). Of note, *Gja1* mRNA was reduced by about 50% in iWAT (Fig. 5F); and this may be related to the very low basal expression of Cx43 in WAT (Fig. 2A).

**Figure 5.**
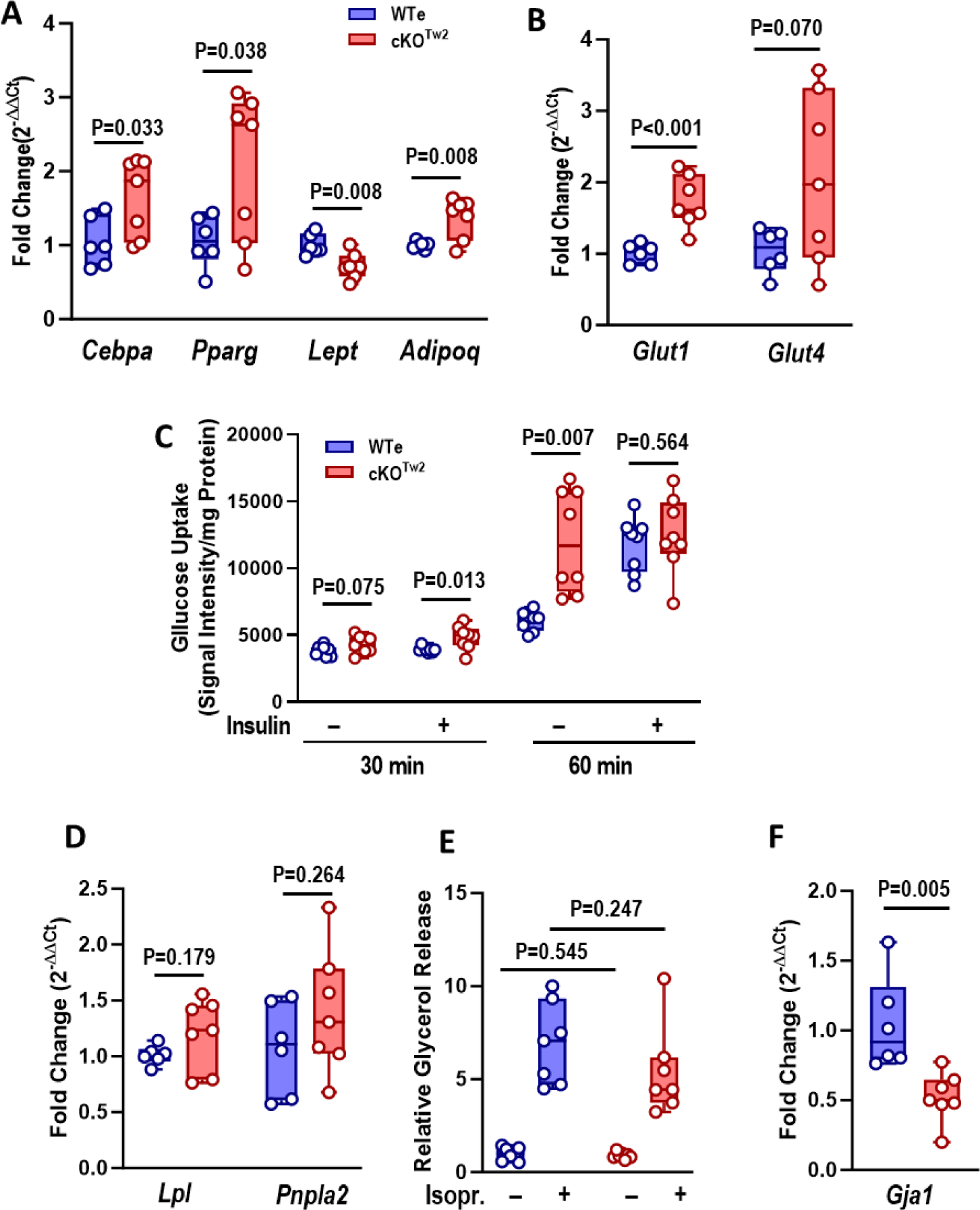
*Gja1* ablation in mesenchymal lineage cells upregulates adipogenic genes and promotes glucose uptake without altering lipolysis in white adipose tissue (WAT). (A) Expression of mRNA by RT-qPCR of adipogenic genes, (B) glucose transporters, (C) 2-NBDG uptake, (D) lipolysis-associated genes, (E) glycerol release relative to tissue weight, in the presence or absence of 10 µM isoproterenol, and (F) *Gja1* mRNA expression in inguinal white adipose tissue of 2-month-old WT (blue) and cKO^TW2^ male mice (red) after 12 weeks on HFD. Boxplots represent interquartile range with median (inside bar); P values were determined by two-sided Mann-Whitney U-test.

Because the *Dermo1/Twist2* promoter targets early mesenchymal precursors, we generated cultures of mesenchymal cells from the ear, which represent adipocyte precursors (18). Consistent with the findings from WAT, adipocytes from cKO^Tw2^ EMSC showed increased expression of *Pparg* and *Adipoq*, as well as *Lep* and the insulin-sensitive glucose transporter *Glut4* (Fig. S5A, B). Such changes were more evident after 5 days in culture, when adipogenic differentiation is more advanced. By contrast, *Cebpa* and *Glut1* mRNA expression did not differ between control and mutant cells (Fig. S5A, B). Confirming effective *Gja1* targeting in these cells (Fig. S5C), *Gja1* mRNA was barely detectable in adipocytes from cKO^Tw2^ EMSC. Overall, these results indicate that lack of Cx43 favors adipogenic differentiation, glucose uptake, and adiponectin production without altering lipolysis.

### *Gja1* ablation in mesenchymal lineage cells increases locomotor activity, food consumption, and energy expenditure

Indirect calorimetry was performed on male mice fed a HFD for 12 weeks. Despite less accumulation of body fat and lower body weight (Fig. 3A, B), cKO^Tw2^ male mice consumed more food compared with WT mice, particularly during periods of activity. The genotype difference was not significant when hourly average consumption was analyzed (Fig. 6A), but it was evident in the average daily food consumed (Fig. 6B). On the other hand, there was a significant genotype effect in hourly energy expenditure, which was higher in cKO^Tw2^ mice during dark cycles (Fig. 6C), although such a difference did not emerge in average daily energy expenditure (Fig. 6D). Since there was a significant interaction between genotype and body weight on energy expenditure (Table S2), these results are consistent with the notion that cKO^Tw2^ male mice consumed more energy per mass unit than their WT littermates, thus offsetting the lower energy consumption expected for their smaller weight. Furthermore, cKO^Tw2^ mice fed a HFD showed higher RER relative to WT mice, in both light and dark cycles (Fig. 6E, F, Table S2). They were also more active during the dark cycle (Fig. 6G. H, Table S2), as noted earlier for mice on standard chow diet (Fig. S3G, H). These findings indicate that cKO^Tw2^ mice are physically and metabolically more active than WT mice, and under obesogenic diet they expend more energy and burn more carbohydrates, particularly during physically active cycles.

**Figure 6.**
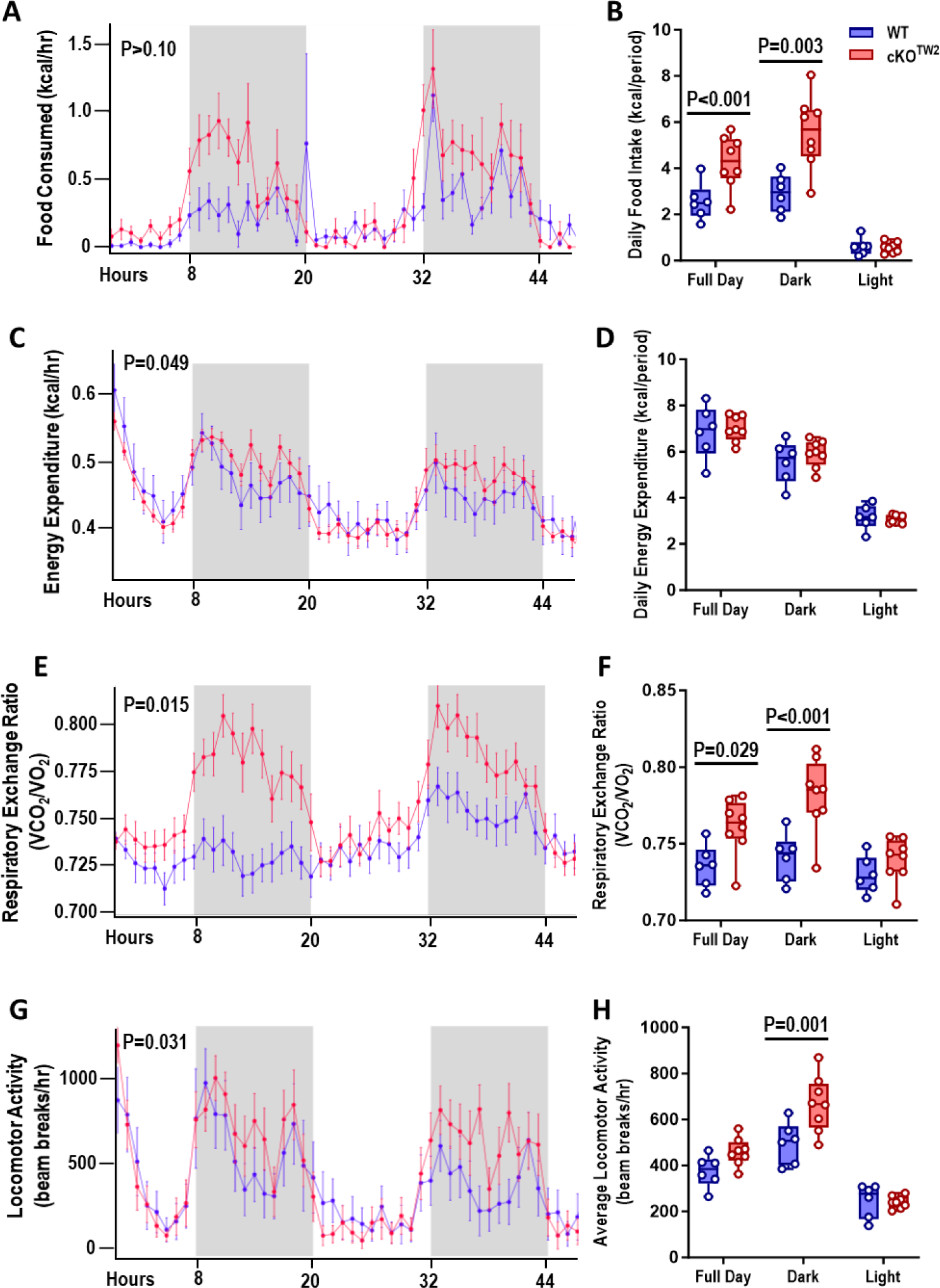
*Gja1* ablation in mesenchymal lineage cells increases locomotor activity, food consumption, and energy expenditure. Two-month-old cKO^TW2^ (red, n=6) and wild type (WT: blue; n=8) male mice were placed in metabolic cages after being fed a high fat diet for 12 weeks, and continuously monitored for 48 hours. (A, B) Food consumption, (C, D) energy expenditures, (E, F) respiratory exchange rate (VCO_2_/O_2_), and (G. H) locomotor activity. Data are presented as both hourly averages (A, C, E, G), analyzed using general linear models or ANOVA (detailed results in Table S2; p values are given for genotype effect), and daily averages over the 2-day experiment for each time period (B, D, F, H), with groups compared using one-way ANOVA.

### *Gja1* ablation in mesenchymal lineage cells protects against obesity-induced BAT whitening and increases cold-induced thermogenesis and lipolysis in mice fed a high fat diet

BAT increases energy metabolism via fatty acid β-oxidation and regulates body temperature through non-shivering thermogenesis (19, 20). Although BAT mass was significantly lower after HFD in cKO^Tw2^ than in WT mice (Fig. 7A, B), histological sections showed substantial fat accumulation in the BAT of WT mice, whereas in cKO^Tw2^ it appeared only minimally infiltrated with white fat cells after HFD (Fig. 7C). Therefore, the larger BAT mass in WT mice reflects BAT whitening under high dietary fat intake, and this change is partially prevented by lack of Cx43. Since Cx43 in BAT is involved in thermoregulation (11), we then studied the response to cold exposure in our mouse model. Both cKO^Tw2^ and WT mice fed a HFD maintained normal body temperature in standard living conditions; however, when exposed to cold (4°C), cKO^Tw2^ mice adjusted to a slightly higher body temperature (about 0.5°C) than WT mice (Fig. 7D). Reflecting higher thermogenesis, we detected higher level of mRNA for *Ucp-1* and for the brown fat-specific genes *Prdm-16* and *Cidea* in BAT of cold-exposed cKO^Tw2^ mice relative to WT mice after HFD (Fig. 7E). As expected, *Gja1* was reduced by 70% in cKO^Tw2^ BAT (Fig. 7E), where non-adipocytic cells expressing Cx43 are present. Consistent with increased BAT activity (19), expression of lipolysis-related genes, *Lpl* and *Pnpla2* was significantly higher in BAT of cKO^Tw2^ mice relative to WT BAT after HFD (Fig. 7F). While ex vivo lipolysis, measured by glycerol release, in resting conditions was not altered in mutant mice, it significantly increased upon β-adrenergic stimulation with isoproterenol in BAT of cKO^Tw2^ mice, but not in WT BAT (Fig. 7G). Finally, BAT of HFD-fed cKO^Tw2^ mice expressed higher mRNA of genes involved in fatty acid oxidation and the OXPHOS system (Fig. 7H, I).

**Figure 7.**
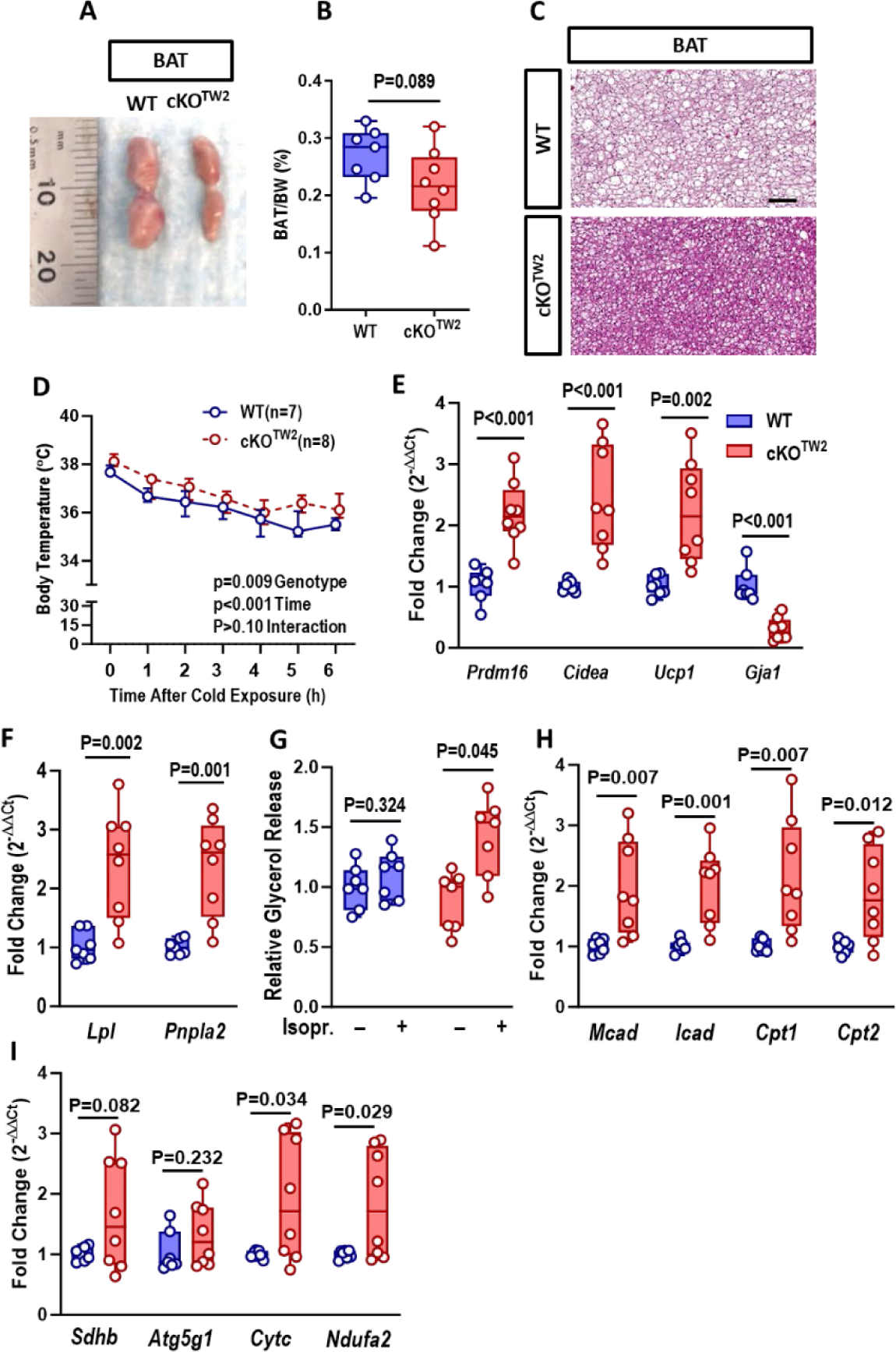
*Gja1* ablation in mesenchymal lineage cells protects from obesity-induced BAT whitening and increases thermogenesis and lipolysis in diet-induced obese mice. (A) Morphology of suprascapular brown adipose tissue (BAT) depots, (B) percent of BAT weight relative to body weight, and (C) representative H&E-stained histological sections of BAT of WT and cKO^TW2^ mice fed a HFD for 12 weeks. Scale bar: 100 µm. Each image is representative of 7 WT and 8 cKO^TW2^ mice. After a 12-week high fat diet (HFD), WT or cKO^TW2^ mice were exposed to cold temperature (4°C) for 6 hours. (D) Core body temperature during the acute cold challenge. Data are shown as average ± 95% CI; P-values represent the effect of genotype, time and their interaction by mixed effects analysis (genotype, F=9.371, p=0.009; time: F=38.74, p<0.001; time x genotype: F=0.6624, p=0.680). Expression of (E) brown adipose tissue (BAT) identity genes, and *Gja1* mRNA, and (F) lipolysis-associated genes in suprascapular BAT depots of 2-month-old WT (blue) and cKO^TW2^ mice (red) after 12 weeks on HFD. (G) Glycerol release relative to tissue weight, in the presence or absence of 10 µM isoproterenol. (H) Expression of β-oxidation and (I) oxidative phosphorylation (OXPHOS) pathway genes in BAT from WT and cKO^TW2^ mice after HFD. Boxplots represent interquartile range and median. Groups were compared using two-sided Mann-Whitney U-test.

### *Gja1* ablation in adipocytes does not protect against diet-induced obesity and actually worsens glucose tolerance

Finally, we asked whether more restricted deletion of *Gja1* in adipocytic cells could reproduce some, if not all, alterations in energy metabolism observed in cKO^Tw2^. To this end, we generated *Adipoq-Cre;Gja1^flox/flox^*(cKO^Adipoq^) mice. As previously described by other groups (11, 12), adipocyte restricted deletion of Cx43 did not protect from the effects of HFD. Unlike cKO^Tw2^ mutants, both male and female cKO^Adipoq^ mice gained as much body weight and fat mass as WT mice after 12 weeks on HFD (Fig. 8A, B; Fig. S6A, B). No differences between controls and mutants were noted in WAT depots (Fig. 8C; Fig. S6D, E). However, BAT mass was higher in cKO^Adipoq^ animals at the end of the HFD period (Fig. 8D; Fig. S6C). In both sexes, mutant and control mice were equally hyperglycemic after HFD, though cKO^Adipoq^ had more severe glucose intolerance relative to WT mice (Fig. 8E; Fig. S6F). Both mutant and WT males showed no changes in blood glucose after insulin (Fig. 8G), whereas females were equally responsive (Fig. S6H). Thus, adipocyte specific *Gja1* ablation does not phenocopy the partial resistance to an obesogenic diet observed with broader *Gja1* ablation in mesenchymal precursors, and it actually worsens glucose tolerance, as previously shown (12).

**Figure 8.**
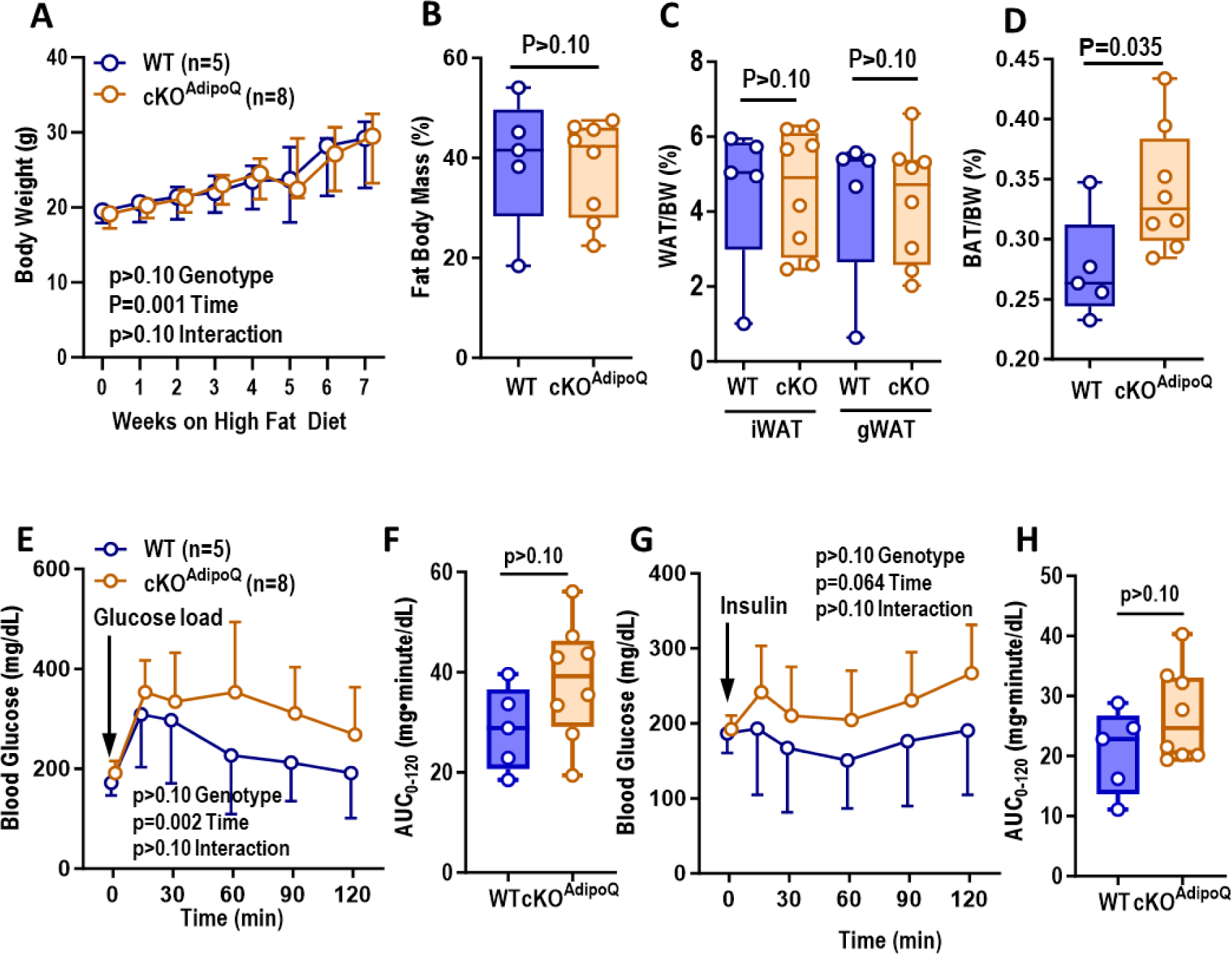
*Gja1* ablation in adipocytic cells does not affect diet-induced obesity and worsens glucose tolerance in mice. (A) Body weight of 2-month-old wild type (WT: blue) and cKO^Adipoq^ male mice (red) during 8 weeks on HFD feeding. Data are shown as median ± interquartile range; P-values represent the effect of genotype, time and their interaction by mixed-effects analysis (genotype, F=0.1452, p=0.710; time: F=51.53, p=0.001; time x genotype: F=0.3383, p=0.9337). (B) Percent body fat by DXA after HFD in the two genotype groups. Percent of (C) inguinal and gonadal white adipose tissue (iWAT, gWAT) and (D) brown adipose tissue depot in the two genotypes after HFD. (E, F) Intraperitoneal glucose tolerance test: blood glucose before and after an intraperitoneal load of 1.5 g/kg glucose. Data points represent the mean ± 95% CI; P-values represent the effect of genotype, time and their interaction by two-way ANOVA (genotype, F=2.603, p=0.135; time: F=6.922, p=0.023; time x genotype: F=0.9072, p=0.483). (G, H ) Intraperitoneal insulin tolerance test: blood glucose before and after an intraperitoneal injection of 0.75U/kg insulin. Data points represent the mean ± 95% CI; P-values represent the effect of genotype, time and their interaction by two-way ANOVA (genotype, F=2.137, p=0.172; time: F=2.822, p=0.064; time x genotype: F=1.014, p=0.418).

## Discussion

We provide evidence that under obesogenic dietary conditions lack of Cx43 in mesenchymal lineage cells leads to increased physical activity, energy expenditure and glucose utilization, reduces WAT fat storage, and mitigates the development of the metabolic syndrome induced by high calorie intake in mice. Therefore, the overall action of Cx43 is to restrain energy consumption and store energy in WAT, and it occurs in both sexes, even though the metabolic advantage of Cx43 ablation is more evident in males than in females. This effect is distinct from Cx43 action in mitochondria, where Cx43 favors energy production and mitochondrial biogenesis (12, 21–23). Thus, Cx43 has lineage- and stage-specific actions that can lead to opposite effects on body adiposity and energy metabolism.

Adipose tissue expansion can occur by either increasing the number of adipocytes (hyperplasia) or increasing the size of existing adipocytes (hypertrophy). Adipocyte number in fat depots is determined early in life and remains rather constant in adult life (24, 25). Accordingly, smaller adipocyte size and lower weight of WAT depots in cKO^Tw2^ mice likely reflects reduced fat hypertrophy in the absence of Cx43 under obesogenic dietary stress. However, we cannot exclude a potential contribution of fat hyperplasia, as HFD upregulated adipogenic genes in *Gja1* ablated WAT and EMSC cultures, and recent evidence suggests that under high caloric stress, adipocyte number can expand around the vasculature of adipose tissues (3, 26). Importantly, smaller WAT adipocytes, increased adipogenic gene expression, reduced BAT whitening, and absence of hepatic steatosis underscores a more metabolically healthy obesity state in cKO^Tw2^ mice (3, 27). Therefore, *Gja1* ablation in *Dermo1/Twist2*-targeted cells reprograms the adipocyte lineage in ways that ultimately enhance the metabolic response to high caloric intake.

Considering its rather broad expression, it is likely that Cx43 acts in multiple mesenchymal-derived cells to drive adipocyte metabolic reprogramming. The enhanced glucose uptake by *Gja1* ablated WAT and EMSC, and increased adipokine expression by EMSC we report here point to a cell autonomous action of Cx43 in adipogenic lineage cells. However, *Gja1* deletion in differentiated adipocytes worsens glucose tolerance without major effects on adiposity under high-calorie stress; and *Adipoq*-driven *Gja1* overexpression increases β-adrenergic stimulated metabolic activity and insulin sensitivity in mice (28). Therefore, adipogenic precursors or cells at early stages of adipogenic differentiation may be key players in the observed phenotype. However, it is likely that other cells targeted by *Dermo1/Twist2* contribute. In an earlier study, we observed modestly lower body weight in mice with *Col1a1*-driven *Gja1* deletion (29); although changes in body mass or composition have not been reported in other models of osteoblast-restricted *Gja1* ablation (14, 30, 31). *Dermo-1/Twist2* also targets muscle (32) and to a lesser extent the liver (this study), two tissues where Cx43 is also expressed, and increased glucose utilization in these tissues may partly explain the observed phenotype. Partial protection from HFD-induced glucose intolerance and insulin resistance, exactly as we report here, and reduced diet-induced inflammation in gonadal adipose tissue has been observed in mice with selective *Gja1* ablation in macrophages (33). As *Dermo-1/Twist2* is expressed by macrophages (34), Cx43 in adipose tissue macrophages may contribute to the phenotype of cKO^Tw2^ mice.

We find that Cx43 is abundantly expressed in adipocyte precursors and rapidly decreases with adipocyte differentiation, as other had also reported (35, 36), while an obesogenic diet upregulates Cx43 expression in BAT. Thus, the function and regulatory mechanisms of Cx43 in the adipose tissue appear to be stage- and tissue-specific. For example, in mature WAT, sympathetic signals upregulate Cx43 to increase body energy metabolism (11); and in mature BAT, Cx43 is required for mitochondrial integrity and increased metabolic activity (12). However, we find that under obesogenic stress, cKO^Tw2^ mice generate more heat after cold exposure, and the BAT of cKO^Tw2^ mice is more metabolically active than the BAT of control mice. Therefore, while the increased physical activity may explain, at least in part, the higher body temperature upon cold exposure, we hypothesize that during embryonic and early postnatal development, lack of Cx43 reprograms BAT precursors towards a more metabolically active phenotype, thus overriding the consequences of lack of Cx43 on mitochondrial function. Our data also suggest that Cx43 in WAT favors fat accumulation and reduces energy expenditure, particularly in obesogenic conditions. The smaller size of fat depots in cKO^Tw2^ mice even under standard dietary regimens may imply that a lower number of adipocytes develop during the early postnatal period, and this may contribute to reduced fat accumulation in obesogenic conditions.

Specific functional domains of Cx43 may have different biologic roles in adipocytes; for example, while chemical inhibition of gap junctional intercellular communication or *Gja1* silencing by siRNA in adipocyte precursors inhibits adipogenic differentiation (7, 35, 37) and prevents WAT beiging in response to cold (11), intercellular communication is dispensable for repression of autophagy by Cx43 (38). Furthermore, Cx43 interacts with the mitochondrial machinery (12) and binds to respiratory complex I (22), actions that do not necessarily require gap junction channel formation. Thus, channel function may be necessary for Cx43 modulation of adipogenesis and the metabolic response to obesogenic diet, while Cx43 action on cellular glucose metabolism and energy production may be linked to channel-independent functions. We also find that, similar to the osteogenic lineage, adipocytes express Cx45, which like Cx43, is downregulated during adipogenic differentiation. These two connexins form gap junctions of different biophysical properties (39), but it is possible that Cx45 may in part compensate for lack of Cx43 in establishing intercellular communication among adipocytes.

The increased locomotor activity certainly contributes to the increased energy expenditure in cKO^Tw2^ mice, particularly under obesogenic diet, the result of increased fuel consumption by muscles and cold exposure while ambulating (40). Notably, while on standard diet cKO^Tw2^ mice use more energy from fat, under obesogenic stress they burn more carbohydrates and utilize more glucose than control mice, particularly during active nocturnal cycles. Such excess energy consumption likely contributes to the lower fat accumulation and better glucose tolerance in cKO^Tw2^ mice on HFD. The reasons for the higher physical activity of mutant mice remain unclear. It is unlikely that this abnormality has a central cause, because the *Dermo1/Twist2* promoter does not target the nervous system, even though Cx43 is present in glial cells (41). Up-regulation of *Adipoq* mRNA in WAT and in differentiating EMSCs supports the idea that increased production of adiponectin or other adipokines, or endocrine factors may in part explain increased glucose utilization and energy consumption by cKO^Tw2^ mice. We and others have shown that Cx43 modulates the expression of factors relevant to bone homeostasis via transcriptional regulation (42, 43), and a similar mechanism might be at play in the adipogenic lineage. It is also possible that adipokines produced by BAT (batokines) may play a role, as Cx43 is abundantly expressed in BAT and is upregulated by high caloric intake.

We propose the following biologic mechanism (Fig. 9): absence of Cx43 in mesenchymal lineages results in reduced volume of white fat depots and reduced WAT hypertrophy and increased glucose utilization by WAT under high caloric intake. In parallel, there is increased fuel utilization, lipolysis and thermogenesis in BAT, and reduced BAT whitening. These changes result in increased energy expenditures, better glucose tolerance, and reduced weight gain and overall better metabolic response to obesogenic stress. Whether increased locomotor activity of Cx43-deficient mice is the main driver of, or it is secondary to a better metabolic balance remains to be determined. We propose that the less severe metabolic syndrome developing in *Gja1*-ablated mice overrides a poorer glucose tolerance caused by loss of Cx43 in the mitochondria of mature adipocytes; thus, limiting the impact of high calorie intake on energy metabolism (Fig. 9).

**Figure 9.**
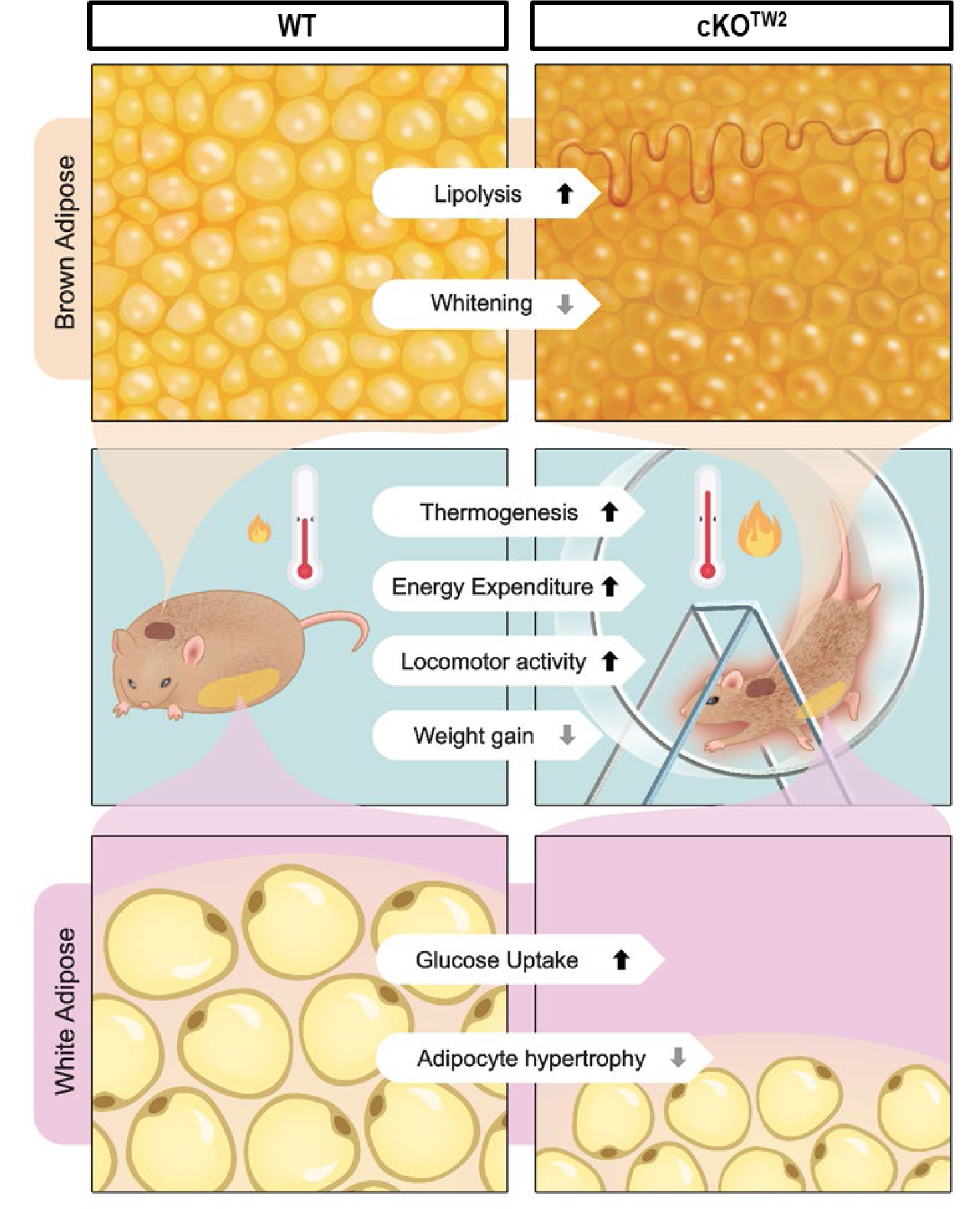
Schematic representation of the effect of *Gja1* ablation on the metabolic response to a high fat diet. (Left column) In normal mice, high dietary calorie intake alters energy metabolism resulting in excess energy storage in fat depots and other organs leading to obesity, hyperinsulinemia, high serum lipids and glucose intolerance. In WAT depots (bottom row), fat accumulation occurs primarily by adipocyte hypertrophy; in BAT (top row), it leads to ‘whitening” as cells become engulfed by lipid droplets. (Right column) Genetic ablation of *Gja1* in the mesenchymal lineage (cKO^Tw2^) results in smaller WAT depots and increased glucose uptake and utilization under high calorie intake. At the organism level (middle row), Cx43-deficient mice are more active and more cold tolerant, burn more energy and utilize more glucose than control littermates under high calorie intake. We propose that the increased energy consumption for physical activity and thermogenesis reduces fat accumulation, WAT hypertrophy, and BAT whitening, resulting in less severe obesity, partially preserved glucose tolerance, and better circulating lipid profile than control littermates.

The increased glucose uptake by cKO^Tw2^ WAT and EMSC suggest that adipose-derived precursors in adult cKO^Tw2^ mice might retain a higher metabolic activity trait in adult animals, and this could be leveraged for therapeutic potential. Infusion of adipose-derived stem cells from human WAT or visceral adipose tissue into obese mice improves glucose metabolism and lipid profiles and reduces body weight gain relative to control mice (44, 45), an effect mediated by secretion of adipokines, anti-inflammatory cytokines, or angiogenic factors (46). Regardless of the mechanism, if this cell-based therapy can be translated to humans, it is conceivable that genetic or pharmacologic interference with Cx43 function in adipose-derived stem cells before infusion in obese subjects may enhance their metabolic activity. This hitherto unknown aspect of Cx43 biology offers a promising new therapeutic target for improving metabolic balance in diabetes and obesity.

## Material and Methods

### Mouse models

The *Dermo1/Tw2-Cre* and *Gja1^flox/flox^* mouse lines and the mating strategy to obtain cKO^Tw2^ mice have been previously described (47). All mice were housed in standard temperature- and humidity-controlled environment with a 12h/12-h light/dark cycle. A diet-induced obesity mouse model was established by feeding high fat diet containing 60% of calories from fat (Research Diet, D12492) for 8 weeks or 12 weeks. After glucose tolerance test and insulin tolerance test (see below), mice underwent dual energy X-ray absorptiometry for body fat composition analysis, followed by fat dissection and collection of brown adipose tissue (BAT) from suprascapular depots, inguinal white adipose tissue (iWAT), and gonadal white adipose tissue (gWAT). For cold exposure experiments, mice fed HFD for 8 weeks were transferred to pre-chilled cages containing pre-chilled water and food in a 4°C cold room. Core body temperature was measured hourly by rectal probe (RET-3, ThermoWorks).

### Blood biochemistry

For intraperitoneal glucose tolerance test, mice fasted for 6 hours were administered a bolus of D-glucose (1.5g/kg) by intraperitoneal injection (IP). For intraperitoneal insulin tolerance test, mice fasted for 6 hours were administered insulin (0.5U/Kg for chow diet-fed mice, and 0.75U/Kg for HFD-fed mice). Blood glucose levels in the tail vein were monitored at various time points (0, 15, 30, 60, 90, and 120 minutes) using the Glucocard Vital blood glucose meter (Arkray, Inc. USA). Serum lipids and hormones were determined by the Washington University Diabetes Models Phenotyping Core, Diabetes Research Center.

### Indirect calorimetry

Respiratory gas measurements, food consumption, and activity were investigated using the TSE/Phenomaster (TSE Systems), also available in the Washington University Diabetes Research Center. Mice were placed in the metabolic cages and monitored for 48 hours. Metabolic variables were elaborated using CalR (https://calrapp.org/). The tool was set to analyze the interaction between body weight (mass effect) and mouse genotype and applies general linear models when a significant interaction exists, or ANCOVA when there is no interaction. ANOVA is used for mass-independent variables (48).

### Real-time quantitative PCR (RT-qPCR)

Total RNA was isolated from fat tissue extracts or EMSCs using RNeasy mini-Kit (Qiagen). Reverse transcription was performed using 500 ng of total RNA and iScript^TM^ reverse transcription super mix (Bio-Rad, cat.1708891). PCR reactions were performed in a 96-well format on an ABI QuantStudio 3 using Fast SYBR Green Master Mix (ABI, cat. 4385612). β2-macroglobulin was used for normalization, and relative expression was calculated by the 2^−(ΔΔCT)^ method. Primer sequences are listed in Table S3.

### Histology

Tissues, fat, and liver were dissected out, fixed in 10% neutral buffed formalin overnight at room temperature, and then were processed for paraffin embedding. Sections (5 µm thickness) were stained with hematoxylin and eosin as previously described (49).

### Immunohistochemistry

Formalin-fixed, paraffin-embedded adipose tissue sections were deparaffinized with xylene, rehydrated and treated with 0.3% hydrogen peroxide in methanol for 15 min to suppress the endogenous peroxidase activity. Antigen retrieval was achieved by placing the samples in a pressure cooker and incubated for 3 min at full pressure in citrate buffer (10 mM citric acid, pH 6.0), followed by gradual cooling to room temperature. Sections were then blocked using serum blocking solution (Invitrogen HISTOSTAIN-SP KIT) and incubated overnight with primary antibody against mouse Cx43 (unconjugated F-7, mouse, Santa Cruz Biotechnology) 1/200 in PBS/0.1% Triton™ X-100 (Sigma-Aldrich) at 4°C. The next day, sections were washed 3 times with PBS and incubated at room temperature with biotinylated universal secondary antibody (Life technologies). After washing with PBS x3, secondary antibodies were visualized using Vectastain ABC (ABC) kit (Vector Laboratories PK-4000) and ImmPACT® DAB Substrate Kit, Peroxidase (HRP) (SK-4105). Sections were then counterstained using Gill II Haematoxylin followed by washing in PBS and dehydration in ascending ethanol series and xylene. Sections were imaged using Hamamatsu NanoZoomer 2.0-HT System.

### Western blotting

As previously described (50, 51), total protein was extracted using RIPA buffer (Cell signaling, cat. 9806) containing protease inhibitor (ThermoFisher Scientific, A32963) and phosphatase inhibitor (ThermoFisher Scientific, A32957). Proteins (15µg) were separated in SDS-PAGE gel by electrophoresis and transferred onto PVDF membrane (Millipore). Membranes were blocked with 5% nonfat dry milk (Cell signaling, 9999) in PBS-T (ThermoFisher, 28352), and blotted using antibodies against Cx43 (Sigma, C6219), Cx45 (Santa Cruz Biotechnology, sc-374354), Cx40 (Santa Cruz Biotechnology, sc-365107), or β-actin (Cell signaling, 4970). Immune reactions were detected using a horseradish peroxidase-conjugated anti-rabbit secondary antibody (Cell signaling, 7074).

### Lipolysis assay

Dissected intrascapular BAT and inguinal adipose tissues were incubated in high-glucose DMEM containing 2% fatty acid-free BSA (Sigma, A8806) for 30 minutes at 37C. To analyze basal lipolysis, tissues were transferred into 96-well plate containing 150µl of high-glucose DMEM supplemented with 2% fatty acid-free BSA and incubated for 1 hr at 37C. To analyze agonist-stimulated lipolysis, tissues were pre-incubated in high-glucose DMEM supplemented with 2% fatty acid-free BSA with 10 µM of isoproterenol for 30 minutes. Tissues were transferred to 96-well plate containing the same medium and incubated for one additional hr. Basal and stimulated lipolysis were determined by measuring glycerol content in the media using free glycerol reagent (Sigma, F6428) and glycerol standard solution (Sigma, G7793). Lipolysis was normalized with protein amount.

### Adipogenic cell culture and differentiation

For ***ear-derived mesenchymal stem cells (EMSC)***, external ears from WT and cKO^TW2^ mouse were collected in ice-cold HBSS containing penicillin/streptomycin (Gibco) and primocin (Invivogen). Ears were cut into small pieces in HBSS containing 2mg/ml of collagenase I (Worthington Biochemical Corporation) and digested for 1h at 37℃ shaking water bath. Digested ears were filtered through 70 µm cell strainer (BD Biosciences) and pelleted by centrifugation at 1300 rpm for 10 minutes. Cells were resuspended using EMSC culture media (DMEM/F12 containing 15% FBS and 10ng/ml of FGF) and seeded in 24-well plates at 2X10^5^ seeding density and incubated for 2 days. For isolation of ***stromal vascular fraction (SVF) pre-adipocytes***, inguinal white adipose tissue was isolated from 2-month-old C57BL/6 mice (Jackson Laboratories) immediately after sacrifice. The tissue was digested by collagenase type I (Sigma) at 37C° for 30 min, and then filtered using a 70µM cell strainer. After centrifugation, the pellet containing the SVF was collected. The cells were resuspended and cultured in DMEM/F12 media (Sigma) with 10% FBS, and the media was changed every other day. To ***induce adipogenic differentiation*** of EMSC, cultures were switched to an adipogenic medium containing 5 µg/ml insulin (Sigma), 1 µM dexamethasone (Sigma), 500 µM IBMX (Sigma), and 5 µM rosiglitazone (Sigma). After 2 days, medium was changed to DMED/F12 containing 10% FBS, 5 µg/ml insulin and 5 µM of rosiglitazone for further 2 days. Then cells were incubated in 10% FBS containing DMEM/F12 until lipid accumulation occurred. For EMSC, cultures were switched to serum-free medium for 2 hr before insulin exposure. The ***ingWAT Mouse Immortalized Preadipocyte Cell Line*** (Millipore cat#: SCC211) was induced to differentiate using AdipoLife DfKt-2 Adipogenesis media (LifeLine Cell Technology), with medium changed every two days. To ascertain adipocyte differentiation, some cultures were stained by Oil Red O (ScienCell) following the manufacturer’s instructions.

### Glucose uptake

Differentiated adipocyte from WT and cKO^TW2^ EMSCs were cultured in growth medium until confluent, then switched to serum free media for 1 hour, before incubation in 100 μM 2-NBD-Glucose (a fluorescent deoxyglucose analog) with or without insulin for 30 or 60 minutes. Fluorescence intensity was measured according to the manufacturer’s instructions (Glucose Uptake Cell-based Assay Kit, Cayman Chemical, 600470).

### Data Presentation and Statistical Analysis

Group data are presented in boxplots with median and interquartile range; whiskers represent maximum and minimum values. Unless otherwise noted, repeated measures are plotted as mean ± 95% confidence interval (CI). Differences between groups were assessed using Mann-Whitney U test, and repeated measures were analyzed by 2-way analysis of variance (ANOVA) or mixed-effects models, in cases of missing data-points, followed by Tukey’s test to adjust p values for multiple comparisons. Data were managed in Microsoft Excel, plotted, and analyzed using Prism 10.0 (GraphPad Software, San Diego, CA).

### Study Approval

All the procedures reported here were approved by the Institutional Animal Care and Use Committee at Washington University (protocol number 20-0029) and followed the Animals in Research: Reporting of In Vivo Experiments (ARRIVE) guidelines.

## Author Contributions

S-Y Lee: designing research studies, conducting experiments, acquiring data, analyzing data, writing the manuscript.

F. Fontana: designing research studies, conducting experiments, acquiring data, writing the manuscript.

T. Sugatani: conducting experiments, acquiring data, writing the manuscript.

I. Portales Castillo: conducting experiments, acquiring data.

G. Leanza: conducting experiments, acquiring data.

Coler-Reilly: analyzing data, generating graphics, writing the manuscript.

R. Civitelli: designing research studies, analyzing data, writing the manuscript, securing funding.

## Acknowledgments

This work was supported by National Institute of Health grant R01 AR041255 and funds from the Barnes-Jewish Hospital Foundation (to RC), by the Alafi Neuroimaging Laboratory, the Hope Center for Neurological Disorders, and NIH Shared Instrumentation Grant (S10 RR0227552) to Washington University, and the Musculoskeletal Research Center (P30 AR074992). Ariella Coler-Reilly was supported by the Skeletal Disorders Training Program (T32 AR060719). The authors wish to thank Dr. Manuela Fortunato, for her preliminary work leading up to the current project, Dr. Maria Remedi, Division of Endocrinology, Metabolism and Lipid Research for her input and guidance, and Dr. David Ornitz, Department of Developmental Biology for guidance on the use of the *Dermo1/Twist2-Cre* mouse model.

## Notes

Conflict of Interest Statement: The authors have declared that no conflict of interest exists

### Competing Interest Statement

The authors have declared no competing interest.

## References

1. Rosen ED, and Spiegelman BM. What We Talk About When We Talk About Fat. Cell. 2014;156(1-2):20–44.

2. Ghaben AL, and Scherer PE. Adipogenesis and metabolic health. Nat Rev Mol Cell Biol. 2019;20(4):242–58.

3. Vishvanath L, and Gupta RK. Contribution of adipogenesis to healthy adipose tissue expansion in obesity. J Clin Invest. 2019;129(10):4022–31.

4. Bartelt A, Bruns OT, Reimer R, Hohenberg H, Ittrich H, Peldschus K, et al. Brown adipose tissue activity controls triglyceride clearance. Nat Med. 2011;17(2):200–5.

5. Berbee JF, Boon MR, Khedoe PP, Bartelt A, Schlein C, Worthmann A, et al. Brown fat activation reduces hypercholesterolaemia and protects from atherosclerosis development. Nat Commun. 2015;6:6356.

6. Schneider-Picard G, Carpentier JL, and Orci L. Quantitative evaluation of gap junctions during development of the brown adipose tissue. J Lipid Res. 1980;21(5):600–7.

7. Yanagiya T, Tanabe A, and Hotta K. Gap-junctional communication is required for mitotic clonal expansion during adipogenesis. Obesity. 2007;15(3):572–82.

8. Burke S, Nagajyothi F, Thi MM, Hanani M, Scherer PE, Tanowitz HB, and Spray DC. Adipocytes in both brown and white adipose tissue of adult mice are functionally connected via gap junctions: implications for Chagas disease. Microbes and infection. 2014;16(11):893–901.

9. Laird DW, and Lampe PD. Therapeutic strategies targeting connexins. Nature reviews Drug discovery. 2018;17(12):905–21.

10. Stains JP, and Civitelli R. Connexins in the skeleton. Semin Cell Dev Biol. 2016;50:31–9.

11. Zhu Y, Gao Y, Tao C, Shao M, Zhao S, Huang W, et al. Connexin 43 Mediates White Adipose Tissue Beiging by Facilitating the Propagation of Sympathetic Neuronal Signals. Cell Metab. 2016;24(3):420–33.

12. Kim SN, Kwon HJ, Im SW, Son YH, Akindehin S, Jung YS, et al. Connexin 43 is required for the maintenance of mitochondrial integrity in brown adipose tissue. Sci Rep. 2017;7(1):7159.

13. Watkins M, Grimston SK, Norris JY, Guillotin B, Shaw A, Beniash E, and Civitelli R. Osteoblast connexin43 modulates skeletal architecture by regulating both arms of bone remodeling. Mol Biol Cell. 2011;22(8):1240–51.

14. Grimston SK, Watkins MP, Stains JP, and Civitelli R. Connexin43 modulates post-natal cortical bone modeling and mechano-responsiveness. BoneKEy Rep. 2013;2:446.

15. Shao M, Wang QA, Song A, Vishvanath L, Busbuso NC, Scherer PE, and Gupta RK. Cellular Origins of Beige Fat Cells Revisited. Diabetes. 2019;68(10):1874–85.

16. Lu X, Altshuler-Keylin S, Wang Q, Chen Y, Henrique Sponton C, Ikeda K, et al. Mitophagy controls beige adipocyte maintenance through a Parkin-dependent and UCP1-independent mechanism. Sci Signal. 2018;11(527).

17. Pettersson US, Walden TB, Carlsson PO, Jansson L, and Phillipson M. Female mice are protected against high-fat diet induced metabolic syndrome and increase the regulatory T cell population in adipose tissue. PloS One. 2012;7(9):e46057.

18. Rim JS, Mynatt RL, and Gawronska-Kozak B. Mesenchymal stem cells from the outer ear: a novel adult stem cell model system for the study of adipogenesis. FASEB J. 2005;19(9):1205–7.

19. Wu Z, Puigserver P, Andersson U, Zhang C, Adelmant G, Mootha V, et al. Mechanisms controlling mitochondrial biogenesis and respiration through the thermogenic coactivator PGC-1. Cell. 1999;98(1):115–24.

20. Lowell BB, and Spiegelman BM. Towards a molecular understanding of adaptive thermogenesis. Nature. 2000;404(6778):652–60.

21. Soetkamp D, Nguyen TT, Menazza S, Hirschhauser C, Hendgen-Cotta UB, Rassaf T, et al. S-nitrosation of mitochondrial connexin 43 regulates mitochondrial function. Basic Res Cardiol. 2014;109(5):433.

22. Boengler K, Ruiz-Meana M, Gent S, Ungefug E, Soetkamp D, Miro-Casas E, et al. Mitochondrial connexin 43 impacts on respiratory complex I activity and mitochondrial oxygen consumption. J Cell Mol Med. 2012;16(8):1649–55.

23. Trudeau K, Muto T, and Roy S. Downregulation of mitochondrial connexin 43 by high glucose triggers mitochondrial shape change and cytochrome C release in retinal endothelial cells. Investigative ophthalmology & visual science. 2012;53(10):6675–81.

24. Hirsch J, and Batchelor B. Adipose tissue cellularity in human obesity. Clin Endocrinol Metab. 1976;5(2):299–311.

25. Spalding KL, Arner E, Westermark PO, Bernard S, Buchholz BA, Bergmann O, et al. Dynamics of fat cell turnover in humans. Nature. 2008;453(7196):783-7.

26. Wang QA, Tao C, Gupta RK, and Scherer PE. Tracking adipogenesis during white adipose tissue development, expansion and regeneration. Nat Med. 2013;19(10):1338–44.

27. Dubois SG, Heilbronn LK, Smith SR, Albu JB, Kelley DE, Ravussin E, and Look AARG. Decreased expression of adipogenic genes in obese subjects with type 2 diabetes. Obesity. 2006;14(9):1543–52.

28. Zhu Y, Li N, Huang M, Chen X, An YA, Li J, et al. Activating Connexin43 gap junctions primes adipose tissue for therapeutic intervention. Acta Pharm Sin B. 2022;12(7):3063–72.

29. Chung DJ, Castro CH, Watkins M, Stains JP, Chung MY, Szejnfeld VL, et al. Low peak bone mass and attenuated anabolic response to parathyroid hormone in mice with an osteoblast-specific deletion of connexin43. J Cell Sci. 2006;119(Pt 20):4187–98.

30. Bivi N, Condon KW, Allen MR, Farlow N, Passeri G, Brun LR, et al. Cell autonomous requirement of connexin 43 for osteocyte survival: consequences for endocortical resorption and periosteal bone formation. J Bone Miner Res. 2012;27(2):374–89.

31. Lloyd SA, Lewis GS, Zhang Y, Paul EM, and Donahue HJ. Connexin 43 deficiency attenuates loss of trabecular bone and prevents suppression of cortical bone formation during unloading. J Bone Miner Res. 2012;27(11):2359–72.

32. Liu N, Garry GA, Li S, Bezprozvannaya S, Sanchez-Ortiz E, Chen B, et al. A Twist2-dependent progenitor cell contributes to adult skeletal muscle. Nat Cell Biol. 2017;19(3):202–13.

33. Choi C, Saha A, An S, Cho YK, Kim H, Noh M, and Lee YH. Macrophage-Specific Connexin 43 Knockout Protects Mice from Obesity-Induced Inflammation and Metabolic Dysfunction. Front Cell Dev Biol. 2022;10:925971.

34. Sun R, Hedl M, and Abraham C. Twist1 and Twist2 Induce Human Macrophage Memory upon Chronic Innate Receptor Treatment by HDAC-Mediated Deacetylation of Cytokine Promoters. J Immunol. 2019;202(11):3297–308.

35. Yeganeh A, Stelmack GL, Fandrich RR, Halayko AJ, Kardami E, and Zahradka P. Connexin 43 phosphorylation and degradation are required for adipogenesis. Biochim Biophys Acta. 2012;1823(10):1731–44.

36. Umezawa A, and Hata J. Expression of gap-junctional protein (connexin 43 or alpha 1 gap junction) is down-regulated at the transcriptional level during adipocyte differentiation of H-1/A marrow stromal cells. Cell Struct Funct. 1992;17(3):177–84.

37. Wiesner M, Berberich O, Hoefner C, Blunk T, and Bauer-Kreisel P. Gap junctional intercellular communication in adipose-derived stromal/stem cells is cell density-dependent and positively impacts adipogenic differentiation. J Cell Physiol. 2018;233(4):3315–29.

38. Bejarano E, Yuste A, Patel B, Stout RF, Jr., Spray DC, and Cuervo AM. Connexins modulate autophagosome biogenesis. Nat Cell Biol. 2014;16(5):401–14.

39. Steinberg TH, Civitelli R, Geist ST, Robertson AJ, Hick E, Veenstra RD, et al. Connexin43 and connexin45 form gap junctions with different molecular permeabilities in osteoblastic cells. EMBO Journal. 1994;13:744–50.

40. Skop V, Guo J, Liu N, Xiao C, Hall KD, Gavrilova O, and Reitman ML. The metabolic cost of physical activity in mice using a physiology-based model of energy expenditure. Molecular metabolism. 2023;71:101699.

41. Penes MC, Li X, and Nagy JI. Expression of zonula occludens-1 (ZO-1) and the transcription factor ZO-1-associated nucleic acid-binding protein (ZONAB)-MsY3 in glial cells and colocalization at oligodendrocyte and astrocyte gap junctions in mouse brain. Eur J Neurosci. 2005;22(2):404–18.

42. Stains JP, Lecanda F, Screen J, Towler DA, and Civitelli R. Gap junctional communication modulates gene transcription by altering the recruitment of Sp1 and Sp3 to connexin-response elements in osteoblast promoters. J Biol Chem. 2003;278(27):24377–87.

43. Gupta A, Leser JM, Gould NR, Buo AM, Moorer MC, and Stains JP. Connexin43 regulates osteoprotegerin expression via ERK1/2 -dependent recruitment of Sp1. Biochem Biophys Res Commun. 2019;509(3):728–33.

44. Liu GY, Liu J, Wang YL, Liu Y, Shao Y, Han Y, et al. Adipose-Derived Mesenchymal Stem Cells Ameliorate Lipid Metabolic Disturbance in Mice. Stem Cells Transl Med. 2016;5(9):1162–70.

45. Calvo E, Keiran N, Nunez-Roa C, Maymo-Masip E, Ejarque M, Sabadell-Basallote J, et al. Effects of stem cells from inducible brown adipose tissue on diet-induced obesity in mice. Sci Rep. 2021;11(1):13923.

46. Sabol RA, Bowles AC, Cote A, Wise R, Pashos N, and Bunnell BA. Therapeutic Potential of Adipose Stem Cells. Adv Exp Med Biol. 2021;1341:15–25.

47. Watkins M, Grimston SK, Norris JY, Guillotin B, Shaw A, Beniash E, and Civitelli R. Osteoblast connexin43 modulates skeletal architecture by regulating both arms of bone remodeling. Mol Biol Cell. 2011;22(8):1240–51.

48. Mina AI, LeClair RA, LeClair KB, Cohen DE, Lantier L, and Banks AS. CalR: A Web-Based Analysis Tool for Indirect Calorimetry Experiments. Cell Metab. 2018;28(4):656–66 e1.

49. Lee SY, Abel ED, and Long FX. Glucose metabolism induced by Bmp signaling is essential for murine skeletal development. Nat Commun. 2018;9.

50. Fontana F, Hickman-Brecks CL, Salazar VS, Revollo L, Abou-Ezzi G, Grimston SK, et al. N-cadherin Regulation of Bone Growth and Homeostasis Is Osteolineage Stage-Specific. J Bone Miner Res. 2017;32(6):1332–42.

51. Revollo L, Kading J, Jeong SY, Li J, Salazar V, Mbalaviele G, and Civitelli R. N-cadherin restrains PTH activation of Lrp6/beta-catenin signaling and osteoanabolic action. J Bone Miner Res. 2015;30(2):274–85.

