## Supplementary Material and Data for "Connexin43 in mesenchymal lineage cells regulates body adiposity and energy metabolism in mice"

### Supplementary Material and Methods

#### Whole embryo mount and X-gal staining

*Tw2-Cre* mice were mated with the *tauLacZ* (Jackson Laboratories; strain 018139) reporter mice, to generate *LacZ:Tw2-Cre* mice. Preparation of whole-body mounts of newborn mice and X-gal staining followed the method previously described (34).

#### Tw2-Cre-mediated cell targeting

*Tw2-Cre* mice were mated with Tm9(CAG-tdTomato)Hze/J (Ai9) mice, to generate Ai9:*Tw2*-Cre mice, and housed as described in Material and Methods. Frozen sections of BAT and WAT were obtained from fresh tissue explanted from 2-month-old mice, immediately after sacrifice, and counterstained with DAPI to visualize nuclei.

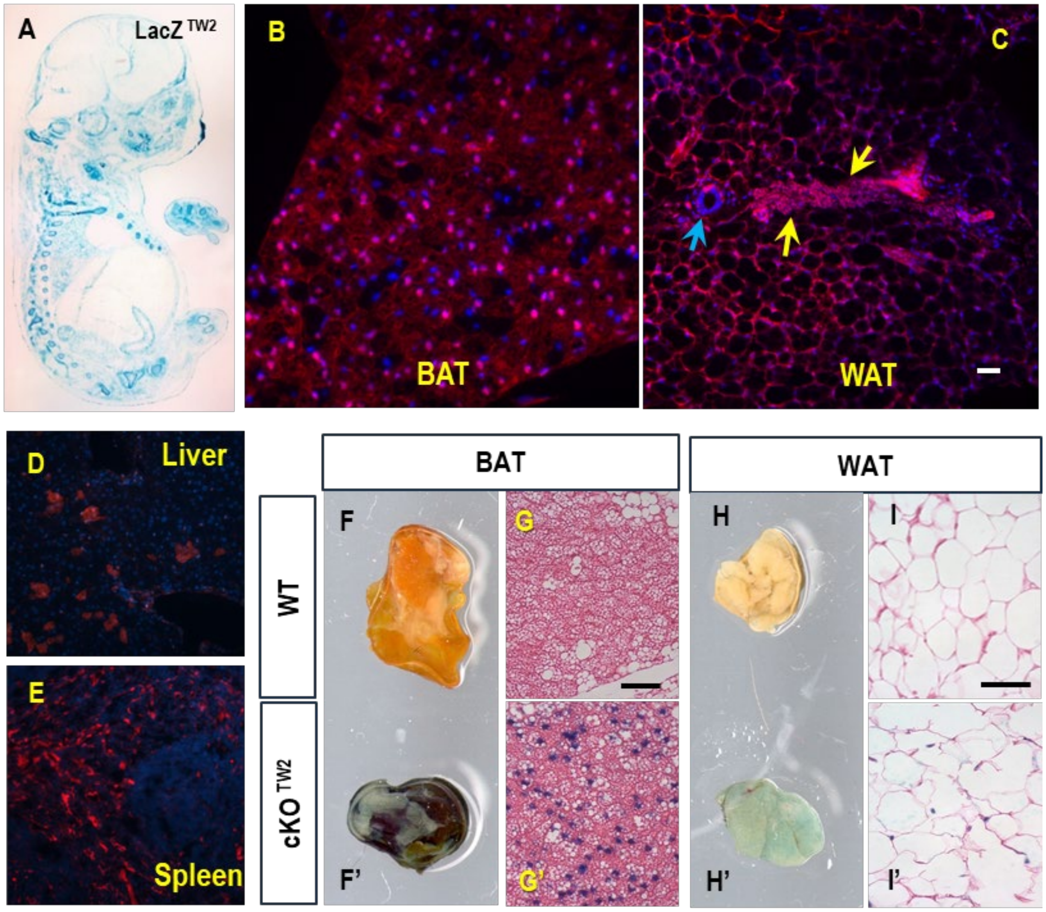


**Supplementary Figure 1. Targeting of adipogenic cells by *Tw2-Cre*.** (A) Whole body mount of a X-gal stained newborn LacZ$^{TW2}$ mouse. (B) Frozen sections of BAT, (C) WAT, (D) liver, and (E) spleen obtained from 2-month-old Ai9:*Tw2*-Cre mice, counterstained with DAPI. Arrows point to red-stained stromal and perivascular cells (yellow) and to the cross-section of a blood vessel, with no red stained cells (cyan). (F) Brown and (H) white adipose tissue (BAT, WAT) isolated from the periscapular area or mammary fat pad, respectively, of 4-month-old cKO$^{Tw2}$ and control littermates (WT) were stained for β-galactosidase (X-gal) activity (dark blue), and counterstained with eosin, before sectioning for (G) BAT and (I) WAT.

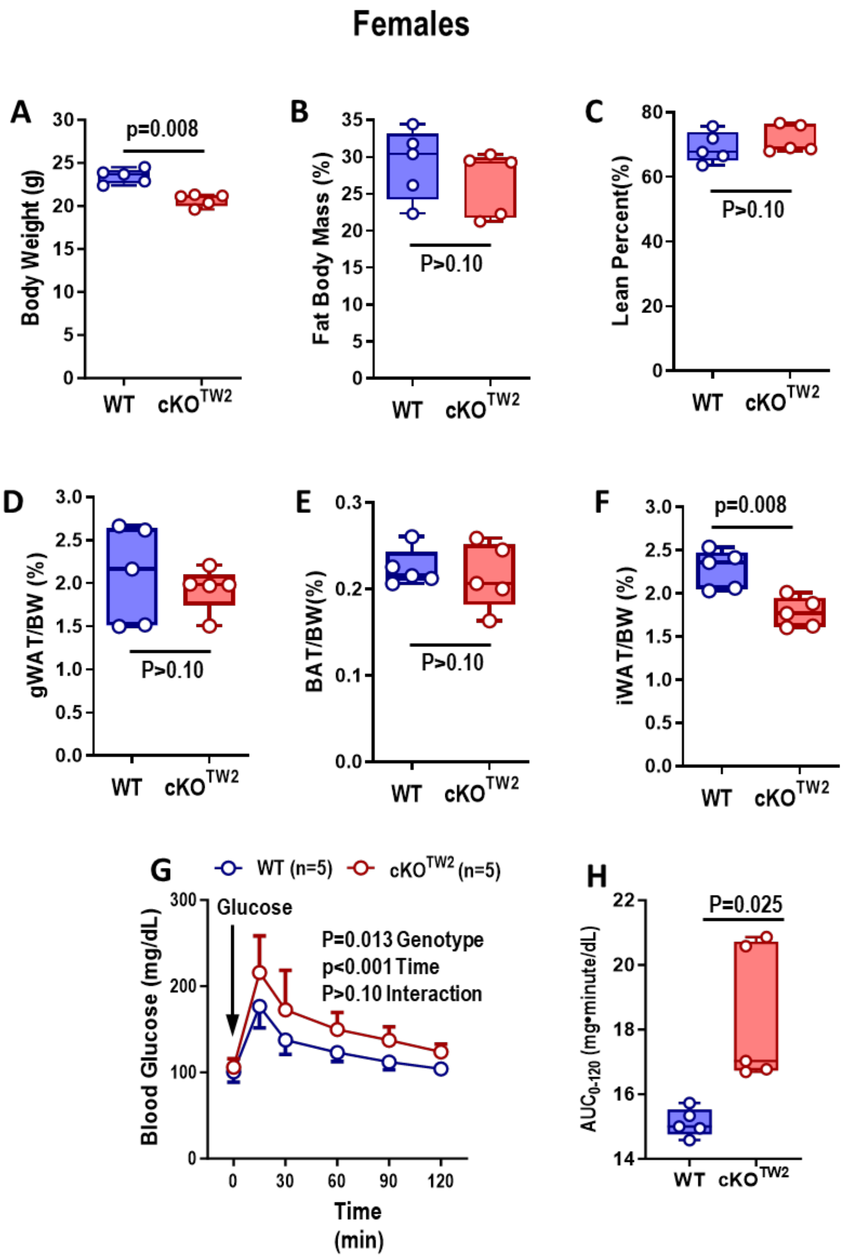


**Supplementary Figure 2.** ***Gja1*** **ablation in mesenchymal lineage cells leads to decreased adiposity in female mice.** (A) Body weight, and (B) DXA-determined percent body fat and (C) lean mass in 7-month-old wild type (WT, blue) $cKO^{TW2}$ female mice (red). (D) Inguinal (iWAT) and (E) gonadal (gWAT) white adipose tissue weight normalized to body weight (BW). (F) Weight of suprascapular brown adipose tissue (BAT) depots normalized to BW. (G) Intraperitoneal glucose tolerance test: blood glucose before and after an intraperitoneal load of 1.5g/kg D-glucose. (H) Areas under the curve (AUC) calculated between 0 and 120 minutes for animals included in panel G. Boxplots represent median and IQR (inside bar). (A-F, H) Groups were compared using two-tailed Mann-Whitney U-test. (G) P-values represent the effect of genotype, time and their interaction by two-way ANOVA (genotype, F=10.11, p=0.013; time: F=56.92, p<0.001; time x genotype: F=1.826, p=0.130).

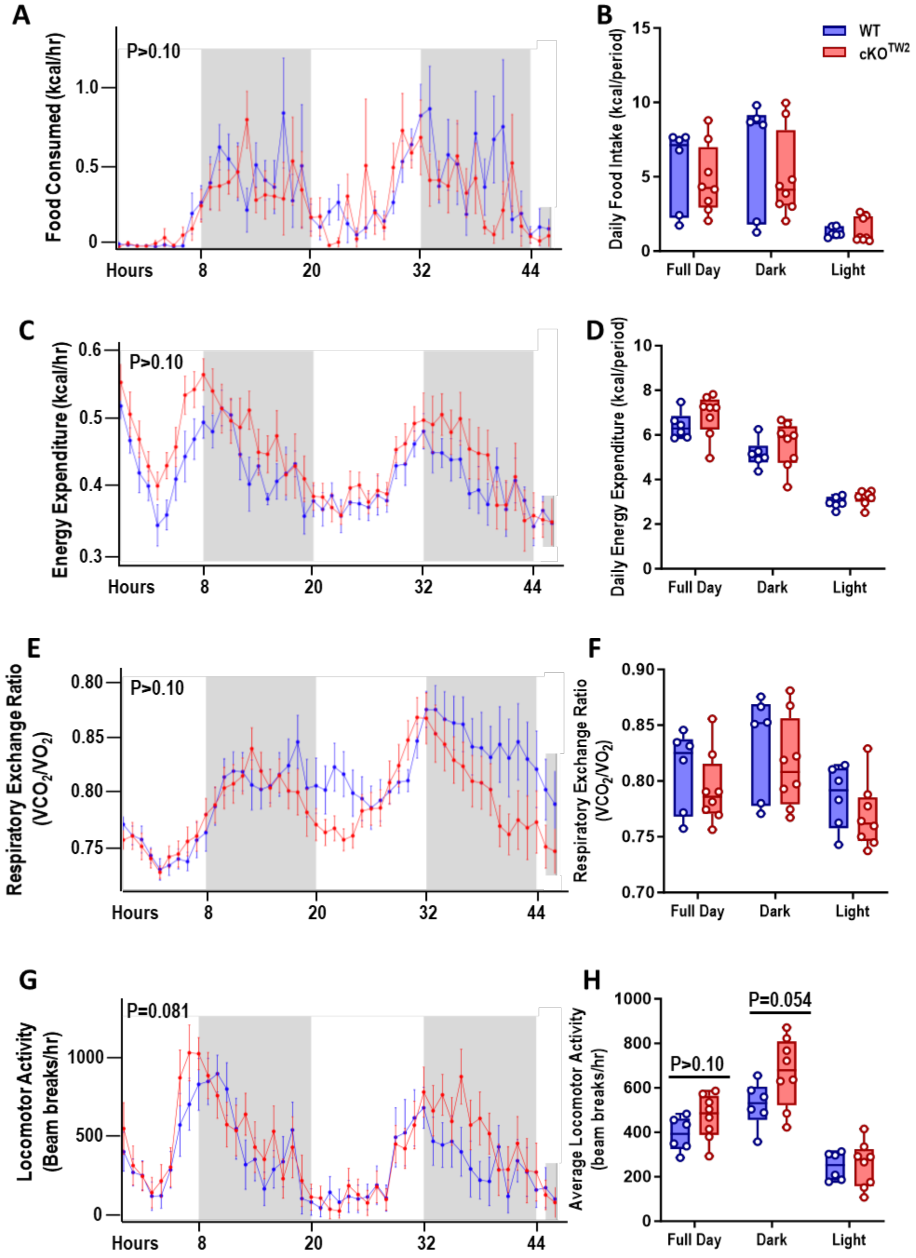


**Supplementary Figure 3. Mice with *Gja1* ablation in mesenchymal lineage cells are more physically active.** Seven-month-old wild type (WT: blue; n=6) and $cKO^{TW2}$ (red, n=8) male mice were placed in metabolic cages, fed a standard diet, and continuously monitored for 48 hours. (A, B) Food consumption, (C, D) energy expenditures, (E, F) respiratory exchange rate ($VCO_2/O_2$), and (G. H) locomotor activity. Data are presented as both hourly averages (A, C, E, G), analyzed using general linear models or ANCOVA (results in Table S1; p values are given for genotype effect), and daily averages for each time period over the 2-day experiment (B, D, F, H), with groups compared using one-way ANOVA.

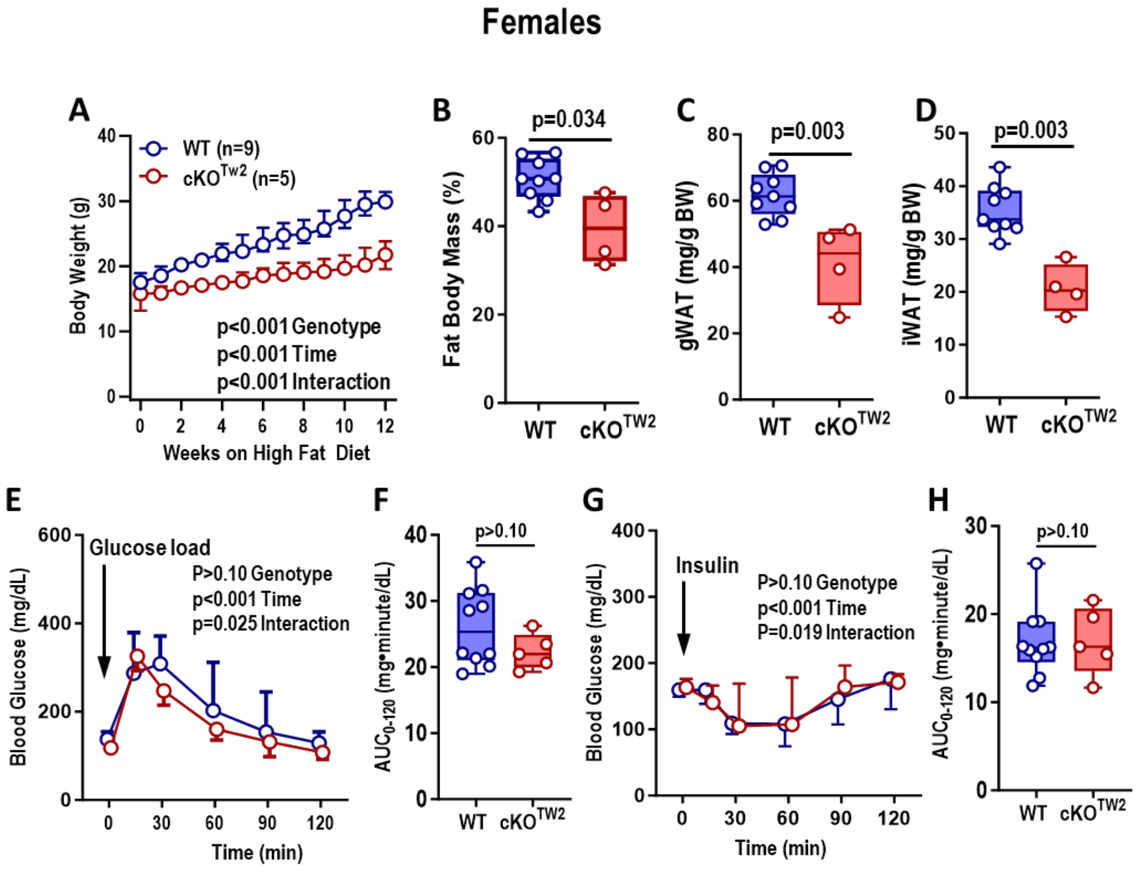


**Supplementary Figure 4. [Formerly part of Fig. 4].** ***Gja1*** **ablation in mesenchymal lineage cells partially protects high fat diet-induced obesity, expansion of fat depots and glucose intolerance in female mice** (A) Body weight of 2-month-old wild type (WT, blue) and $cKO^{TW2}$ (red) female mice during 12 weeks on HFD feeding. Data are shown as median and interquartile range (mixed-effects analysis: genotype, F=38.09, p<0.001; time: F=127.8, p<0.001; time x genotype: F=13.75, p<0.001). (B) Percent body fat, gWAT (C) and iWAT (D) mass in the two genotype groups. (E) Intraperitoneal glucose tolerance test: blood glucose before and after an intraperitoneal load of 1.5g/kg D-glucose (mean ± 95%CI; two-way ANOVA: genotype, F=1.488, p=0.244; time: F=78.37, p<0.001; time x genotype: F=2.771, p=0.025). (F) Area under the curve (AUC) calculated between 0 and 120 minutes for animals included in panel E. (G) Intraperitoneal insulin tolerance test: blood glucose before and after an intraperitoneal injection of 0.75U/kg insulin (mean ± 95%CI; two-way ANOVA: genotype, F=0.038, p=0.8497; time: F=13.60, p<0.001; time x genotype: F=2.920, p=0.019). (F) Area under the curve (AUC) calculated between 0 and 120 minutes for animals included in panel G. Group data are in boxplots representing the interquartile range with median (inside bar), and P values were determined by two-sided Mann-Whitney U-test.

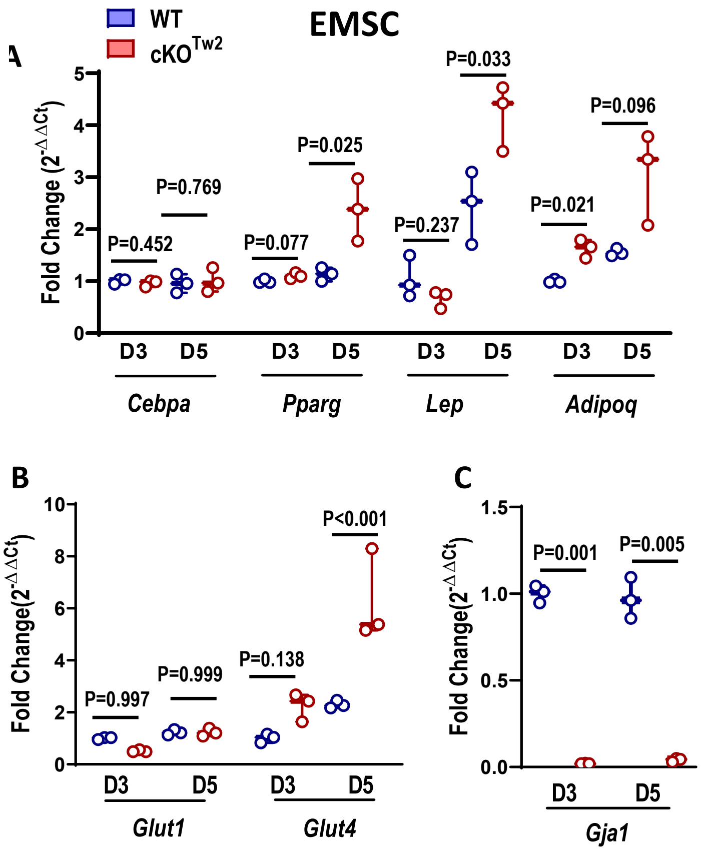


**Supplementary Figure 5. [Formerly Sup Fig 2] Increased expression of adipokines and glucose transporters in *Gja1*-deficient ear-derived mesenchymal stem cells (EMSC)** (A) Expression of mRNA by RT-qPCR of adipogenic genes, (B) glucose transporters, and (C) *Gja1* mRNA in EMSC cultured from either WT (blue) or $cKO^{TW2}$ (red) male mice after 12 weeks on HFD. Boxplots represent median and IQR (inside bar), and P values were determined by two-sided Mann-Whitney U-test.

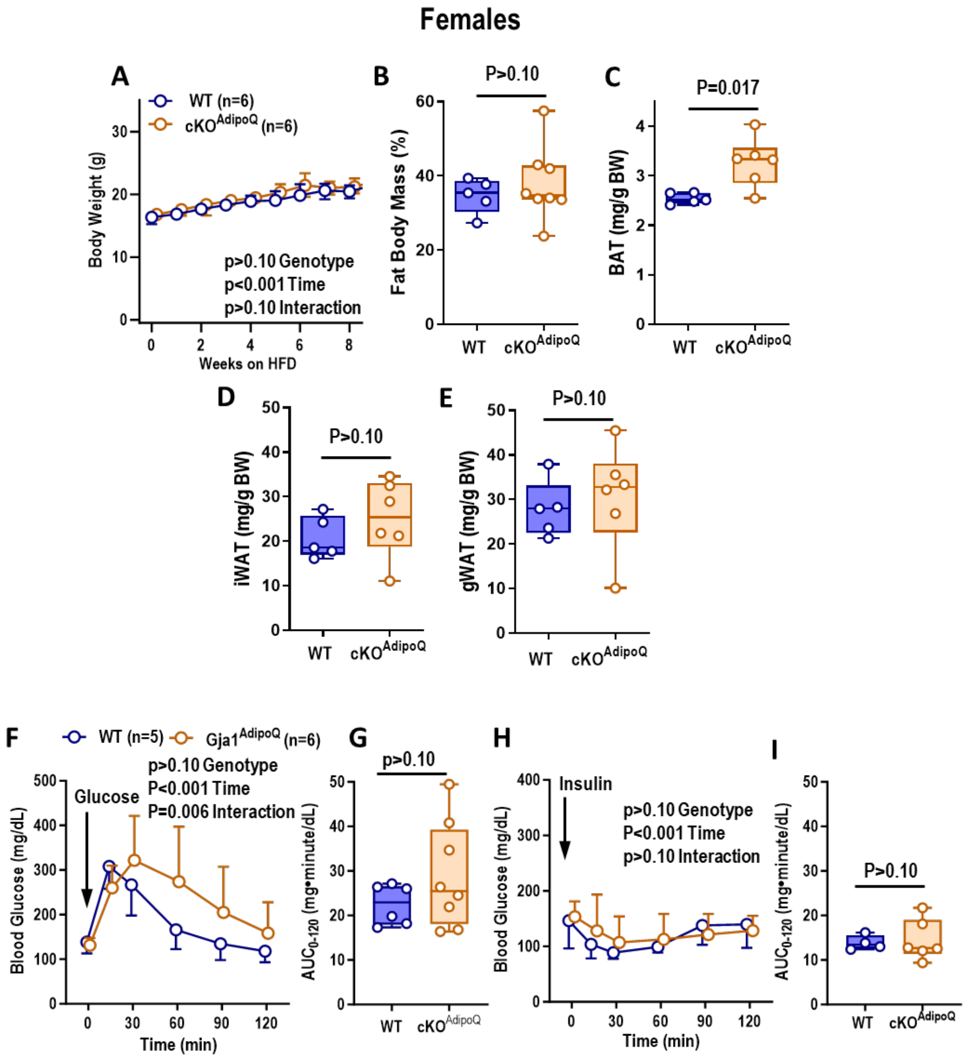


**Supplementary Figure 6. [Formerly Sup Fig 3] *Gja1* ablation in adipocytic cells does not affect diet-induced obesity and worsens glucose tolerance in female mice.** (A) Body weight of 2-month-old wild type (WT: blue) and $cKO^{Adipoq}$ female mice (orange) during 9 weeks on HFD feeding. Data are shown as median ± IQR; P-values represent the effect of genotype, time and their interaction by two-way ANOVA (genotype, F=1.442, p=0.257; time: F=95.77, p<0.001; time x genotype: F=0.6861, p>0.10). (B) Percent body fat by DXA after HFD in the two genotype groups. (C) Brown adipose tissue (BAT) weight; (D) Inguinal (iWAT) and (E) gonadal (gWAT) white adipose tissue weight relative to body weight after HFD. (F) Intraperitoneal glucose tolerance test: blood glucose before and after an intraperitoneal load of 1.5 g/kg glucose (mean ± 95% CI; two-way ANOVA: genotype, F=1.056, p=0.324; time: F=23.46, p<0.001; time x genotype: F=3.667, p=0.006). (G) Areas under the curve (AUC) calculated between 0 and 120 minutes for animals included in panel F. (H) Intraperitoneal insulin tolerance test: blood glucose before and after an intraperitoneal injection of 0.75U/kg insulin (mean ± 95% CI; two-way ANOVA: genotype, F=0.093, p=0.768; time: F=9.592, p<0.001; time x genotype: F=1.716, p=0.153). (I) Areas under the curve (AUC) calculated between 0 and 120 minutes for animals included in panel H. Boxplots represent the interquartile range with median (inside bar), and P values were determined by two-sided Mann-Whitney U-test.

Table S1. Statistical Analysis of Indirect Calorimetry Data for Male Mice on Standard Chow Diet

| General Linear Models (p values) | | | | | | | | | |
|---|---|---|---|---|---|---|---|---|---|
| | Full Day | | | Light | | | Dark | | |
| | Weight | Genotype | Interaction | Weight | Genotype | Interaction | Weight | Genotype | Interaction |
| *Food Consumed (kcal/period)* | **0.050** | 0.956 | >0.10 | **0.043** | 0.334 | >0.10 | 0.131 | 0.737 | >0.10 |
| *Water Consumed (ml/period)* | 0.107 | 0.562 | >0.10 | 0.355 | 0.425 | >0.10 | **0.080** | 0.809 | >0.10 |
| *Energy Expenditure (kcal/period)* | 0.427 | 0.245 | >0.10 | 0.646 | 0.211 | >0.10 | 0.343 | 0.299 | >0.10 |
| *Oxygen Consumption (ml/hr)* | 0.483 | 0.229 | >0.10 | 0.736 | 0.203 | >0.10 | 0.377 | 0.277 | >0.10 |
| *Carbon Dioxide Production (ml/hr)* | 0.266 | 0.328 | >0.10 | 0.354 | 0.265 | >0.10 | 0.247 | 0.393 | >0.10 |

| ANOVA (p values for genotype effect) | | | |
|---|---|---|---|
| | Full Day | Light | Dark |
| *Respiratory Exchange Ratio* | 0.375 | 0.422 | 0.395 |
| *Locomotor Activity (beam breaks)* | 0.133 | 0.434 | 0.186 |
| *Ambulatory Activity (beam breaks)* | **0.081** | 0.206 | 0.170 |

Shown are p-values obtained by applying general linear models using body weight as covariate, and one-way ANOVA for mass-independent variables (analyzed by CalR). Data were collected over two, consecutive full day cycles (48 hours). Highlighted in bold are p-values lower than 0.10.

Table S2. Statistical Analysis of Indirect Calorimetry Data for Male Mice Kept on High Fat Diet for 8 Weeks

| General Linear Models (p values) | | | | | | | | | |
|---|---|---|---|---|---|---|---|---|---|
| | Full Day | | | Light | | | Dark | | |
| Effect | Weight | Genotype | Interaction | Weight | Genotype | Interaction | Weight | Genotype | Interaction |
| *Food Consumed (kcal/period)* | 0.502 | 0.319 | >0.10 | 0.227 | 0.246 | >0.10 | 0.679 | 0.136 | >0.10 |
| *Water Consumed (ml/period)* | 0.724 | 0.363 | >0.10 | 0.705 | 0.400 | >0.10 | 0.789 | 0.734 | >0.10 |
| *Energy Expenditure (kcal/period)* | 0.114 | **0.049** | **0.047** | 0.564 | 0.450 | >0.10 | 0.144 | **0.053** | **0.041** |
| *Oxygen Consumption (ml/hr)* | 0.119 | **0.049** | **0.048** | 0.583 | 0.454 | >0.10 | 0.151 | **0.055** | **0.043** |
| *Carbon Dioxide Production (ml/hr)* | 0.098 | **0.052** | **0.047** | 0.477 | 0.427 | >0.10 | 0.126 | **0.057** | **0.041** |

| ANOVA (p values for genotype effect) | | | |
|---|---|---|---|
| | Full Day | Light | Dark |
| *Respiratory Exchange Ratio* | **0.015** | 0.203 | **0.004** |
| *Locomotor Activity (beam breaks)* | **0.031** | 0.416 | **0.004** |
| *Ambulatory Activity (beam breaks)* | **0.062** | 0.424 | **0.010** |

Shown are p-values obtained by applying general linear models using body weight as covariate, and one-way ANOVA for mass-independent variables (analyzed by CalR). Data were collected over two, consecutive full day cycles (48 hours). Highlighted in bold are p-values lower than 0.10.

Table S3. Primers used for RT-qPCR

| | Forward | Reverse |
|---|---|---|
| *AdipoQ* | TGTTCCTCTTAATCCTGCCCA | CCAACCTGCACAAGTTCCCTT |
| *Atg5g1* | GCTGCTGAGAGATGGGTTC | AGTTGGTGTGGCTGGATCA |
| *Atgl* | GCCACTCACATCTACGGAGC | GACAGCCACGGATGGTGTTC |
| *β2M* | TCACATGTCTCGATCCCAGT | GGGAAGCCGAACATACTGAA |
| *Cebpα* | CAAGAACAGCAACGAGTACCG | GTCACTGGTCAACTCCAGCAC |
| *Cidea* | ATCACAACTGGCCTGGTTACG | TACTACCCGGTGTCCATTTCT |
| *Cpt1* | CCCATGTGCTCCTACCAGAT | CCTTGAAGAAGCGACCTTTG |
| *Cpt2* | AGCCAGTTCAGGAAGACAGA | GACAGAGTCTCGAGCAGTTA |
| *Cytc* | TCCATCAGGGTATCCTCTCC | GGAGGCAAGCATAAGACTGG |
| *Flk1* | GGGATGGTCCTTGCATCAGAA | ACTGGTAGCCACTGGTCTGGTTG |
| *Gja1* | CGGTTGTGAAAATGTCTGCTATG | GGCACAGACACGAATATGATCT G |
| *Gja4* | CCCACATCCGATACTGGGTG | CGAAGACGACCGTCCTCTG |
| *Gjc1* | AGATCCACAACCATTCGACATTT | TCCCAGGTACATCACAGAGGG |
| *Glut1* | TACGTGGAGCCCTAGGCACA | TGTAGCAGGGCTGGGATGAA |
| *Glut4* | AAAAGTGCCTGAAACCAGAG | TCACCTCCTGCTCTAAAAGG |
| *iCad* | GTAGCTTATGAATGTGTGCAACTC | GTCTTGCGATCAGCTCTTTCATTA |
| *Leptin* | GAGACCCCTGTGTCGGTTC | CTGCGTGTGTGAAATGTCATTG |
| *Lpl* | CAGCTGGGCCTAACTTTGAG | GACCCCCTGGTAAATGTGTG |
| *mCad* | GGCCATTAAGACCAAAGCAGA | GTGTCGGCTTCCACAATGAAT |
| *Ndufa2* | GCACACATTTCCCCACACTG | CCCAACCTGCCCATTCTGAT |
| *Pparγ* | TCGCTGATGCACTGCCTATG | GAGAGGTCCACAGAGCTGATT |
| *Prdm16* | CAGCACGGTGAAGCCATTC | GCGTGCATCCGCTTGTG |
| *Sdhb* | GGACCTATGGTGTTGGATGC | GTGTGCACGCCAGAGTATTG |
| *Ucp1* | AGGCTTCCAGTACCATTAGGT | CTGAGTGAGGCAAAGCTGATTT |
